# Method for Extracellular Electrochemical Impedance Spectroscopy on Epithelia

**DOI:** 10.1101/2025.04.02.646694

**Authors:** Athena Chien, Colby F. Lewallen, Hanna Khor, Analia Vazquez Cegla, Rongming Guo, Nael A. McCarty, Kapil Bharti, Craig R. Forest

## Abstract

Epithelial tissues form barriers to the flow of ions, nutrients, waste products, bacteria, and viruses. The conventional electrophysiology measurement of transepithelial resistance (TER) can quantify epithelial barrier integrity, but does not capture all the electrical behavior of the tissue or provide insight into membrane specific properties. Electrochemical impedance spectroscopy, in addition to measurement of TER, enables measurement of transepithelial capacitance (TEC) and a ratio of electrical time constants for the tissue, which we term membrane ratio. This protocol describes how to perform galvanostatic electrochemical impedance spectroscopy on epithelia using commercially available cell culture inserts and chambers, detailing the apparatus, electrical signal, fitting techniques, and error quantification. The measurement can be performed in approximately one minute using instrumentation capable of galvanostatic sinusoidal signal processing (4 *μ*A amplitude, 2 Hz-50 kHz). All fits to the model have less than 10 Ω mean absolute error, revealing repeatable values distinct for each cell type. On representative retinal pigment (n=3) and bronchiolar epithelial samples (n=4), we measured TER 500-667 Ω.cm^2^ and 955-1034 Ω.cm^2^, within the expected range, TEC 3.65-4.10 *μ*F*/*cm^2^ and 1.07-1.10 *μ*F*/*cm^2^, and membrane ratios 18-22 and 1.9-2.2, respectively.

## 1 Introduction

Epithelial electrophysiology can provide insight into epithelial barrier function by measuring the electrical properties associated with the transport of ions, nutrients, and waste products in an environment similar to in vivo conditions [1, 2]. Further, transport across the tissue can be perturbed by blocking and activating ion channels or degrading tight junctions formed between the cells to isolate specific pathways or channels. This perturbation is useful for understanding healthy and diseased models, testing therapeutics, and characterizing quality control of in vitro cultures [3–5].

Transepithelial resistance (TER, also known as transepithelial electrical resistance, TEER) has become the gold standard over the last 30 years for quantifying epithelial tissue maturity and barrier integrity using commercial or custom electrodes and chambers [1, 6, 7]. Electrochemical impedance spectroscopy (EIS/ECIS), although not yet widely adopted, has been shown to offer insights into epithelial transport dynamics, membrane-specific properties, and more accurate measurement of membrane integrity. For example, Lewallen et al. showed the advantages of combining EIS with intracellular voltage recording to distinguish apical and basolateral transport function during adenosine triphosphate administration [3] in a custom Ussing chamber and automated electrode placement [8]. Cottrill et al. reports similar parameters without intracellular voltage recording, by making assumptions about relative permeability [9]. Linz et al. integrates EIS into existing commercial electrodes (STX, World Precision Instruments) to measure Caco-2 monolayer dynamics in response to stimuli: saponin, sonoporation, and calcium chelator ethyleneglycol-bis(beta-aminoethylether)-N,N’-tetraacetic acid (EGTA) [10].

We describe a protocol to perform galvanostatic EIS on commercially available cell culture inserts (3460 Membrane Insert, Corning) within electrophysiology chambers (EndOhm, World Precision Instruments) with guidelines on the apparatus, frequencies, fitting technique, and error quantification. Using a frequency sweep from 2 Hz to 50 kHz, the 4 *μ*A sinusoidal waveform results in the measurement of TER, TEC, and membrane ratio in less than one minute. All fits to the model have less than 10 Ω mean absolute error, revealing repeatable values distinct for each cell type. On representative retinal pigment (n=3) and bronchiolar epithelial samples (n=4), we measured TER measurements 500-667 Ω.cm^2^ and 955-1034 Ω.cm^2^, TEC measurements 3.65-4.10 *μ*F*/*cm^2^ and 1.07-1.10 *μ*F*/*cm^2^, and membrane ratio measurements 18-22 and 1.9-2.2, respectively.

## 2 Motivation

Transepithelial resistance (TER) is a good indicator of tight junction development and membrane integrity. Tight junctions form between the cells [1, 7, 10], creating the paracellular pathway as shown in Fig. 1A. The electrical resistance of the tissue is dominated, but not determined exclusively, by this paracellular pathway, compared to the flow of ions through the respective apical and basolateral membranes.

**Figure 1.**
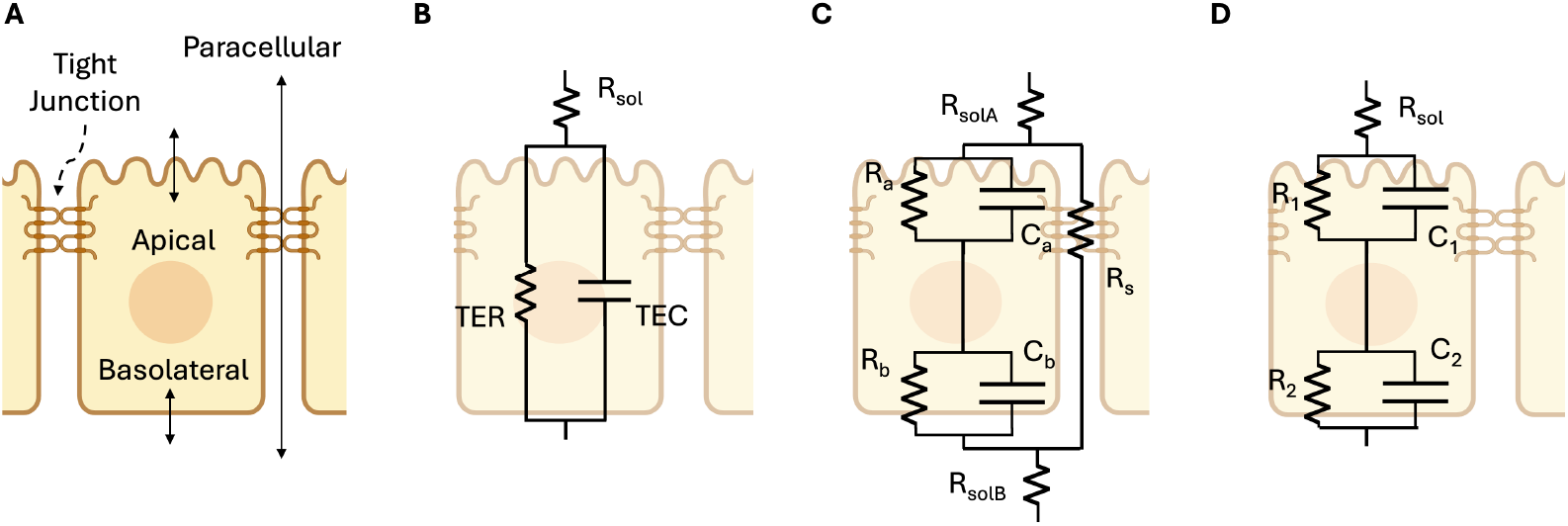
Circuit models for transport across an epithelial monolayer modified from Lewallen et al.[3] (A) Physical model: cross-section schematic of an epithelial tissue showing the three pathways where transport is regulated: apical, basolateral, and paracellular pathways. (B) 3-parameter circuit model: a simple resistor-capacitor (RC) circuit that can be fit from extracellular electrodes alone, widely used to measure trans-epithelial resistance (TER) and trans-epithelial capacitance (TEC) despite its low fidelity. (C) 7 parameter model with apical, basolateral and paracellular pathways requires intracellular voltage measurement to fully solve [3, 11]. (D) RCRC model, shown to model the apical and basolateral cell membranes of Caco-2 cells in Linz et al. [10]

TER measurement is typically performed at a single frequency (e.g., 12.5 Hz or 75 Hz [7, 12]) assuming resistance is independent of frequency (i.e., reactance is negligible). Yet it has been observed that reactance is not negligible [3, 10, 13], and therefore this assumption is not valid. As a result, the TER measured with a signle frequency is erroneously low. For example, bronchiolar epithelia show a strong frequency dependent change in resistance (See Fig. 2). Mathematical models of epithelia that include reactance contributed by capacitance, as shown in Fig. 1B-D, can more accurately fit the measurements across frequencies. Fig. 2 shows an EIS data fit with model from Fig. 1D. In this figure, called a Nyquist plot, the resistance (R) and reactance (X) are the real and imaginary components of the complex impedance (Z), measured as a function of frequency.

**Figure 2.**
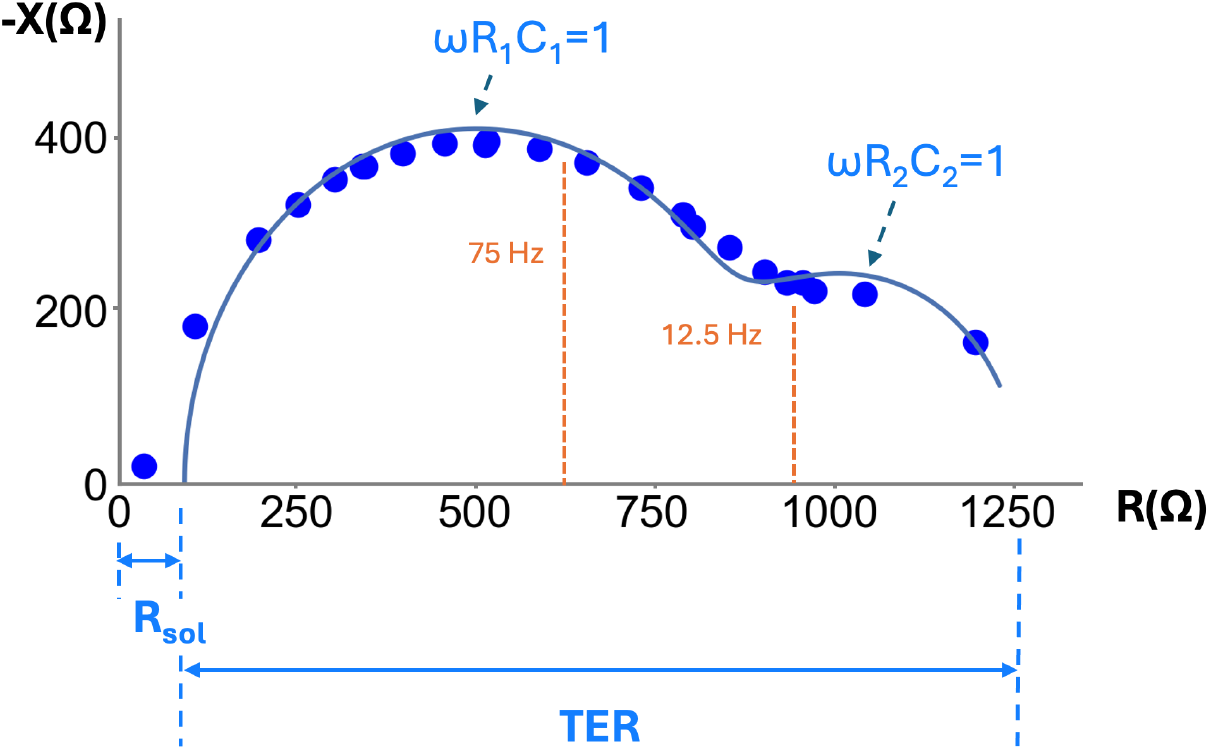
Nyquist plot of negative reactance (-X) vs. resistance (R) for frequencies 2 Hz - 10 kHz on a measurement of bronchiolar epithelia (at day 7). Each point is the measured reactance and resistance for a given frequency, all measurements were taken on a single sample. TER measurements taken at a single frequency, such as 12.5 Hz and 75 Hz, underestimate TER [18] and require background subtraction. Each capacitance component can be estimated based on the approximate peak of each semicircular curve in the Nyquist plot, where at the peak, 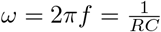.

Adding capacitive elements to the model (Fig. 1B-D) can yield novel insights about the epithelial barrier. EIS measurements of TER from fitting a circuit model, as shown in Fig. 2, inherently separates electrical resistance of the tissue from the transwell and media, termed *R*_solution_, shortened to *R*_sol_. Thus unlike traditional TER measurement, “background subtraction” from a blank is therefore not necessary. The impedance measurements also capture capacitance, enabling one to model charge blockage or storage at the tissue surface, a frequency dependent property [3, 10, 13]. In addition, transepithelial capacitance (TEC) shows promise in describing cell properties such as cell volume, surface area, and even health and vitality [14–17]. Although cell capacitance has been used for single cells to identify cell types in cytometry, there has yet to be rigorous biological studies published correlating the TEC of a monolayer of cells to a specific cell property.

Fig. 1B is the simplest model that accounts for capacitance, and it lumps the apical and basal membrane, along with the paracellular properties, into single values of TER and TEC. While insightful and implemented in commercial devices (CellZscope, NanoAnalytics), we have shown that some epithelial tissues, like bronchiolar epithelia, require more detailed models to accurately calculate TER (e.g. Fig. 1C-D [19]). Fig. 1C, while modeling the apical, basolateral, and paracellular pathways separately, requires intracellular measurement to solve [3], thus the method we present here relies on the model of Fig. 1D. This model can capture the different time constants observed in the data (*τ* = *RC*), yet can be measured with only extracellular electrodes. This model has been used in prior publications to analyze impedance data on Caco-2 cells changing in response to forskolin [10]. One of the shortcomings, however of the model of Fig. 1D is that the paracellular resistance is lumped into *R*_1_, *R*_2_, *C*_1_, *C*_2_ and therefore not resolvable.

Thus, with off-the shelf electronic circuitry and software for fitting an electronic model of epithelia that we have developed, one can measure membrane specific properties. The hardware relies on the same transwells and chambers, with their integrated extracellular electrodes, as are used for conventional TER measurement. The electronic circuitry enables galvanostatic signal processing (e.g,. analog to digital conversion, modulation) and conversion to impedance. The custom software performs the math and data analysis (e.g., fitting and error calculation).

The electrical current is set to less than 10 *μA* to maintain linearity [13, 14], and the frequency sweep is chosen such that the reactance is approximately zero at the bounds of the range, or approximately 2 Hz-50 kHz, with frequencies logarithmically spaced.

Prior to describing the method for epithelia, we will first discuss our validation of the EIS and fitting technique on electrical circuits that mimic the TER and TEC of living tissues. We soldered circuits with commonly observed TER and TEC values of mature tissues, and lower TER and TEC values representing developing tissues, resulting in four electrical circuits: 16HBE1, 16HBE2, RPE1, and RPE2 shown in Fig. 3. The results show precise and accurate TER (< 7Ω error, <1 Ω SD), TEC (< 0.12*μ*F error (< 0.025*μ*F SD)), R_sol_(< 4Ω error (< 0.4Ω SD)) and membrane ratio < 1.5 error (< 0.6 SD) for all four circuit models. Individual circuit parameters *R*_sol_, *R*_1_, *R*_2_, *C*_1_, *C*_2_ and their errors are reported in Supplemental Figure 15.

**Figure 3.**
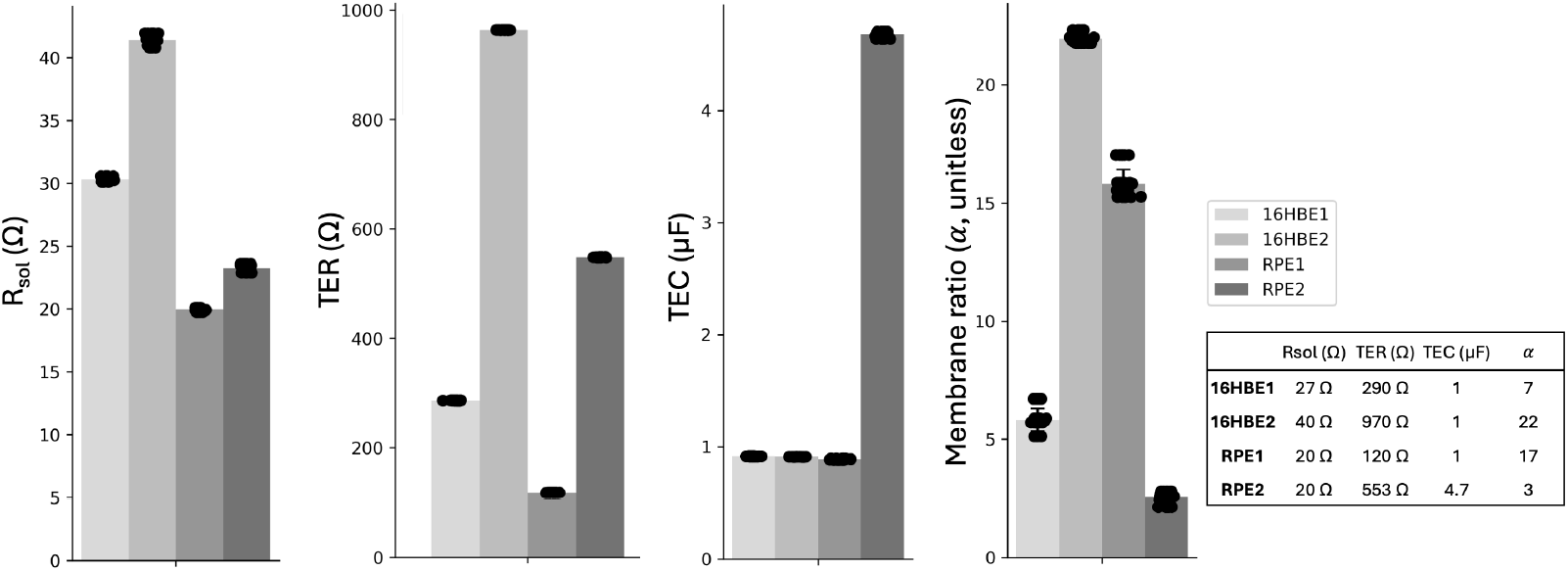
EIS measurement and fitting to RCRC model (Fig. 1D) on four electrical circuits (16HBE1, 16HBE2, RPE1, RPE2) simulating 16HBE and RPE cell layers. Six replicate EIS measurements taken on each circuit, and all six measurements were fit six times to include fitting variability. TER (< 7Ω error, <1 Ω SD), TEC (< 0.12*μ*F error (< 0.025*μ*F SD)), R_sol_(< 4Ω error (< 0.4Ω SD)) and membrane ratio < 1.5 error (< 0.6 SD) for all 4 circuit models. Signal 2 Hz - 50 kHz, 2 freq/dec, 4 *μ*A. Table inset: actual circuit values.

## 3 Materials and Reagents

### 3.1 Cells

For Human Bronchiolar Epithelia:

1. 16HBE14o- Human Bronchiolar Epithelial Cells (Millipore Sigma, catalog number: SCC150) grown to confluency (7 days after seeding at 250k cells/12-well transwell in 12 mm Transwell® with 0.4 μm Pore Polyester Membrane Insert (Corning, catalog number: 3460))
2. 16HBE Culture Media (concentrations based on medium used by Cystic Fibrosis Foundation [20]):
  a. 89% MEM Media (Thermo Fisher Scientific, catalog number: 11095098)
  b. 10% Fetal Bovine Serum, Premium, Heat Inactivated (R&D Systems, S11150H)
  c. 1% Penicillin-Streptomycin (5,000 U/mL) (Thermo Fisher Scientific, catalog number: 15070063)

For Retinal Pigment Epithelia:

1. iPSC-derived Retinal Pigment Epithelia grown to maturity (Sharma et al. 2022 [21])) in 12 mm Transwell® with 0.4 μm Pore Polyester Membrane Insert (Corning, catalog number: 3460)
2. RPE Cell Culture Media (Sharma et al. 2022 [21])

### 3.2 Biological supplies

1. 1x 12 mm Transwell® with 0.4 μm Pore Polyester Membrane Insert with no cells seeded (Corning, catalog number: 3460)
2. Spray bottle (Fisher Scientific, catalog number: 02-991-721)
3. 2x 50 mL Falcon tube (Corning, catalog number: 352070)
4. Reagent-grade Alcohol (VWR, catalog number: BDH1164-4LP)
5. Lint-free wipes (VWR, catalog number: 89218-057)
6. Deionized water (e.g., Millipore Milli-Q Water Purification System)
7. Sodium Hypochlorite, 5% w/v (LabChem, catalog number: 1310-73-2)
8. P1000 pipette (VWR, catalog number: 83009-776)
9. P1000 tips (VWR, catalog number: 76322-154)
10. Sterile Forceps (VWR, catalog number: 82027-408)

### 3.3 Electrical supplies

1. EndOhm (12mm, World Precision Instruments, catalog number: EndOhm-12G)
2. EndOhm to RJ11 Cable (World Precision Instruments, catalog number: 53330-01)
3. Autolab banana cable (Metrohm)
4. Autolab power cable (Metrohm)
5. USB-A to USB-B 2.0 cable (Metrohm)
6. USB-C to USB-A adapter (j5Create, catalog number: JCD383)
7. Autolab test cell (Metrohm, catalog number: 3500002470)
8. Parts for RJ11 to Banana adapter
9. 5x Banana plug (Digikey, catalog number: J356-ND)
  a. RJ11 Female Connector (Amazon Uxcell, catalog number: B01HU7BX42)
  b. Plastic Housing Box (P200 tip box, VWR, catalog number: 76322-150)
  c. M12 Drill Bit (Drill America, catalog number: POUM12X1.25)
  d. Solder (Digikey, catalog number: SMDIN52SN48-ND)
  e. Superglue (Loctite, catalog number: 1799408)

## 4 Equipment

1. Multi Autolab cabinet for M204 modules (Metrohm, catalog number: AUT.MAC204.S)
  a. M204 multichannel PGSTAT module (Metrohm, catalog number: AUTM204.S)
  b. FRA32M module for M101/204/PGSTAT204 (Metrohm, catalog number: FRA32M.MAC.204.S)
2. PC with CPU 1 GHz or faster 64 bit processor, 2 GB RAM, 20 GB hard disk space, DirectX 9.0c compliant display adaptor with 64 MB RAM (Dell Laptop, CPU intel i9 core, 2.50 GHz, 64 GB RAM, 1.84 TB Hard drive, intel(R) UHD graphics GPU)
3. Laminar flow hood/biological safety cabinet (Labconco, Logic+ Type A2 Biosafety Cabinet)
4. Soldering iron (Digikey, catalog number: PES51-ND)
5. Drill (Dewalt, catalog number: DCD793B)
6. Multimeter with continuity feature (Fluke, catalog number: FLUKE-177 ESFP)

## 5 Software

1. Matlab 2024b
2. Microsoft Excel (Version 16.92)
3. NOVA 2.1 (Metrohm, 2024)
4. Impedance analysis github (extracellularEIS, https://github.com/chien0507/extracellularEIS/tree/main)
  a. Matlab impedance fitting code (Nova batch 20240125.m, Nova function 20240125.m, fitZ12 20240125.m, funRCRC.m, makeTissueID.m)
  b. NOVA custom galvanostatic EIS protocol (protocol.nox)

## 6 Procedure

### 6.1 Fabricate Adapter

1. Fabricate “RJ11 to Banana Adapter” according to Fig. 4 as follows: using the drill and drill bit, drill holes into the top of the plastic housing box, taking care not to crack the housing, for the five banana plugs (labeled GND, WE, S, CE, and RE). Insert plugs and secure them using the associated metal nuts. Also drill one hole for the RJ11 adapter and glue in place with superglue. Solder each wire on the RJ11 Female Connector to a metal tab on the back of a banana plug, there should be one banana plug that is not soldered to anything that serves as our floating ground (GND). Connect the EndOhm to the RJ11 to banana adapter. Determine connections using the multimeter continuity feature by touching one probe to the banana plug in question and the other probe to each of the corresponding EndOhm electrodes labelled in Fig. 4. Label the banana plugs accordingly (RE for Reference, S for Sense, CE for Counter, WE for working).

**Figure 4.**
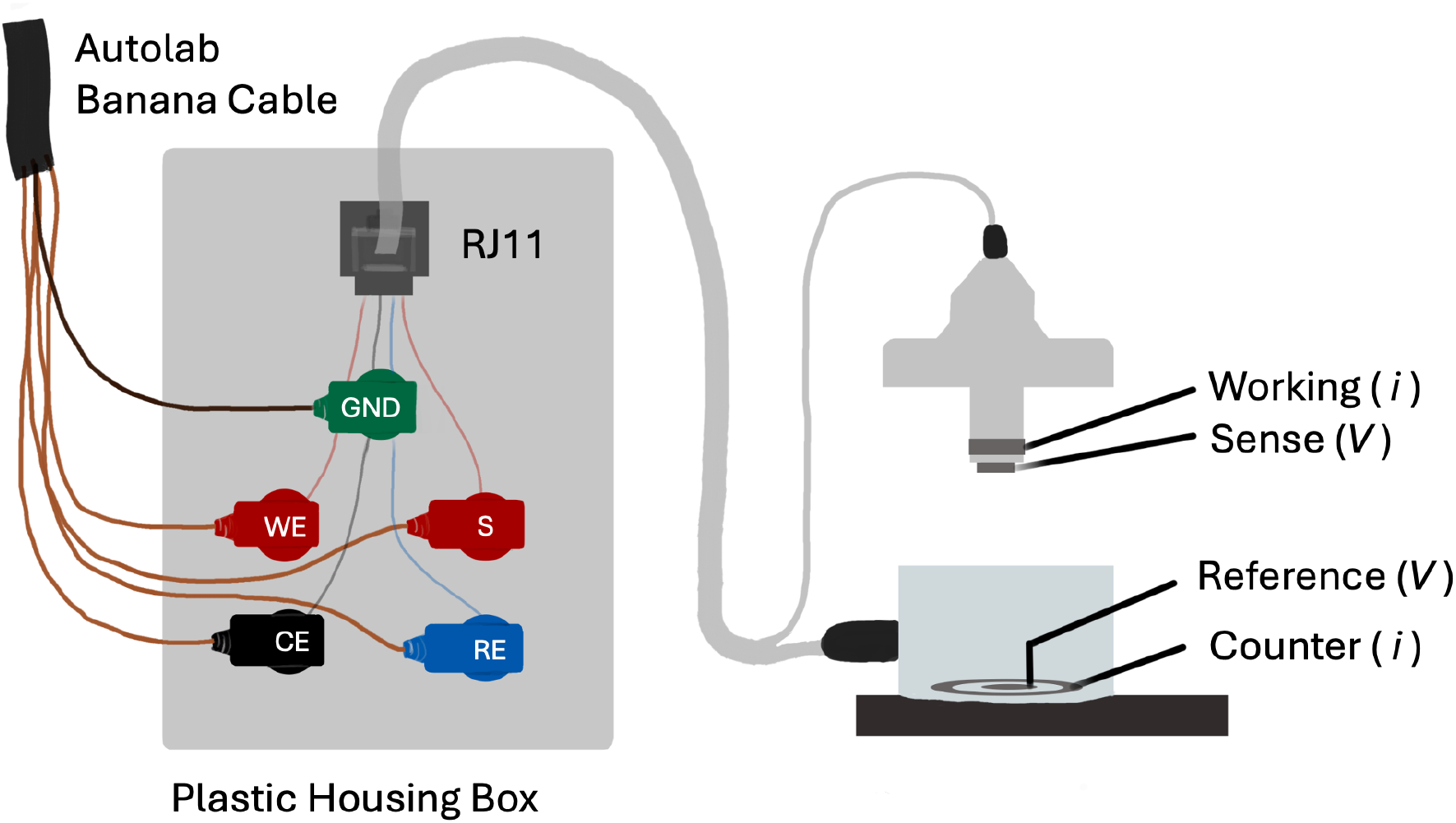
The RJ11 to Banana Adapter enables one to connect the Autolab M204 to the EndOhm. In performing the EIS measurement, the electrical current is sent across the working and counter electrodes and the resulting voltage is measured across the reference and sense electrodes.

### 6.2 Experimental Setup

1. Fill spray bottle with reagent-grade alcohol.
2. Fill 50 mL Falcon tube with ∼ 20 mL alcohol and place forcep tips in alcohol. Close top and shake gently to sterilize entire forceps. Place Falcon tube into sanitized laminar flow hood for transferring cell culture inserts later.
3. In the laminar flow hood, transfer 5 mL of media (specific to which cell line is being used) to the remaining Falcon tube using the P1000 pipette and tips. Wait 10 minutes for the temperature of media in Falcon tube to reach room temperature.
4. Check EndOhm electrodes for any nicks or scratches, or lightness in color that indicate rechloriding is necessary. If so, follow the instructions to rechloride [22]. Briefly, fill the EndOhm with sodium hypochlorite solution. Let the electrodes soak for 10 minutes. Electrode should appear darker in color. Rinse with deionized water.

### 6.3 Control Impedance Measurements

This section describes how to take control measurements on the Autolab test cell and on a blank transwell to ensure the system is connected and working properly prior to starting biological measurements. When measuring on the blank transwell and for all measurements on epithelia, the system will be set up as shown in Fig. 5.

**Figure 5.**
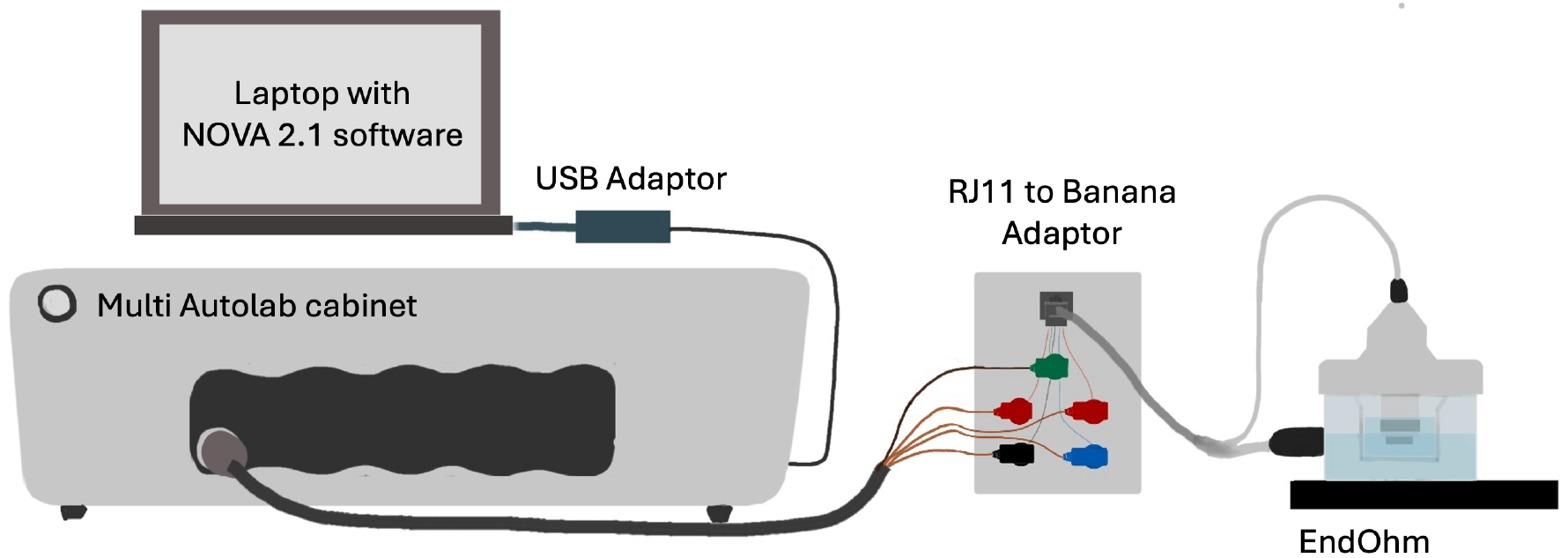
Diagram of instrument connections for extracellular impedance spectroscopy. The galvanostat is connected to a chamber containing a transwell and electrodes for measurement.

1. Spray two lint-free wipes with reagent alcohol and set inside the flow hood. Sterilize the EndOhm by spraying both the interior and exterior of the EndOhm generously with reagent alcohol before setting it to dry on the lint-free wipes for at least 10 minutes.
2. While waiting for the EndOhm to dry, plug the following into the Multi Autolab cabinet: Autolab banana cable, USB-A to USB-B 2.0 cable, and power cable connected to wall outlet. Turn on the Multi Autolab cabinet. Connect the USB-B 2.0 to the PC (using the USB-C to USB-A adapter if necessary).
3. Connect the Autolab banana cable to the Autolab test cell, matching up the cable ends as shown in Fig. 7(left). This connects the Autolab through the test cell’s 1 kΩ and 1 *μ*F parallel RC circuit.
4. On the PC, open the NOVA software, open the Github link and download the repository by clicking the green “code” button and “Download ZIP.” Find the downloaded zip folder in File Explorer and double click the compressed file. Then select “extract all” to unzip the file. The NOVA procedure to use is in the unzipped file, titled “protocol.nox.”
5. In the NOVA software home screen, click “Import Procedure” and open the “protocol.nox.” file. The protocol sends a galvanostatic signal at 4 *μ*A with 2 frequencies per decade, 1 second wait time before starting the measurement, and 5 frequencies stacked at low frequencies (5sines). If more datapoints are desired for each sweep, you can increase the frequencies per decade.
6. As shown in Fig. 6, in the opened protocol, select the “Repeat n times” block, then select the “FRA measurement” block, and finally the “Freq Export” block.
7. In the properties side panel that opens, enter the filename including the file path. For the filepath, we recommend naming the file in the form “mediaUsed TW1 day1 exp1 freq” but any filename without spaces will work. Impedance data is exported as a tab delimited text file. This protocol outputs the resistance and reactance (real and imaginary impedance) for each frequency and the transepithelial voltage measured just before each frequency sweep.
8. On the NOVA software, start the measurement by clicking “Measurement” in the top menu, “Run on” and then the channel the test cell is plugged into. A warning will pop up saying “current range (10 *μA*) is not sufficient for selected frequency (50000 Hz).” Click “OK.” Once the measurement begins, the software switches to a new open tab on the left with the collected data. The Nyquist plot should appear in the bottom row of plots. Verify that the Nyquist plot generated resembles the shape shown in Fig. 7. This data will be analyzed later to verify that the model cell resistance and capacitance are accurately measured. Data will automatically be exported at the end of the measurement. Close the data tab.
9. Disconnect the M204 banana cables from the test cell, and connect them to the EndOhm using the RJ11 to banana adapter. Spray and wipe down the EndOhm to RJ11 cable, before bringing the EndOhm end into the laminar flow hood and connecting to the EndOhm as shown in Fig. 4. The adapter and M204 remain outside of the flow hood, so temporarily taping EndOhm to RJ11 cable to the edge of the laminar flow hood prevents excess strain on the cable.
10. Next we will measure the impedance of a blank transwell, to verify that the blank transwell’s electrical properties are accurately measured. Depending on which cells will be measured in subsequent steps, add 2.5 mL of the corresponding cell culture media to the EndOhm using the P1000 pipette and tips. Avoid inducing bubbles by pipetting slowly.
11. In the laminar flow hood, take out the sterile tweezers in reagent alcohol and shake lightly to dry off. Using the tweezers, place the blank or empty cell culture insert into the EndOhm bottom. To prevent air bubbles getting trapped on the bottom of the cell culture insert surface, contact the media surface at an angle before placing it level and releasing. Bubbles can be dislodged by lightly shaking the cell culture insert in the media solution, or removing and replacing. See Fig. 8 for what bubbles may look like on the bottom of the cell culture insert.
12. Pipette 700 *μ*L of media into the apical side of the cell culture insert, and place the EndOhm cap on top, once again lowering one edge in first to prevent bubbles. Bubbles tend to get stuck on the side of the apical electrodes, and can be removed by pipetting the apical media over the electrode surface first, or by brushing any bubbles on the edge of the EndOhm bottom after the initial dip. If bubbles appear, lift and reset the cap, or pipette media towards the bubbles to dislodge and pop them from the bottom of the EndOhm. If bubbles are present, they may contribute deleteriously to impedance measured. Examples of bubbles present on an EndOhm electrode are shown in Fig. 8.
13. Change the filename and filepath in the NOVA protocol again to reference the blank cell culture insert, as described in steps 6 and 7 and shown in Fig. 6. For the filename, we recommend naming the file in the form “mediaUsed TW1 day1 exp1 freq” but any filename without spaces will work.
14. Run measurement as in step 8 on the blank cell culture insert. Also note that the reactance should be distributed around a mean of zero as shown in Fig. 9A. If the Nyquist plot shows a nearly vertical line, either the electrodes are not properly submerged in media, the cable is not connected, or the device is not running on the correct channel (see Fig. 9B). This blank measurement is not necessary for every experiment, but is useful to check the system is connected and working properly. Notably, bubbles will result in a larger scatter of real and imaginary impedances (Fig. 9A).
15. Remove the blank cell culture insert and close the data tab.

**Figure 6.**
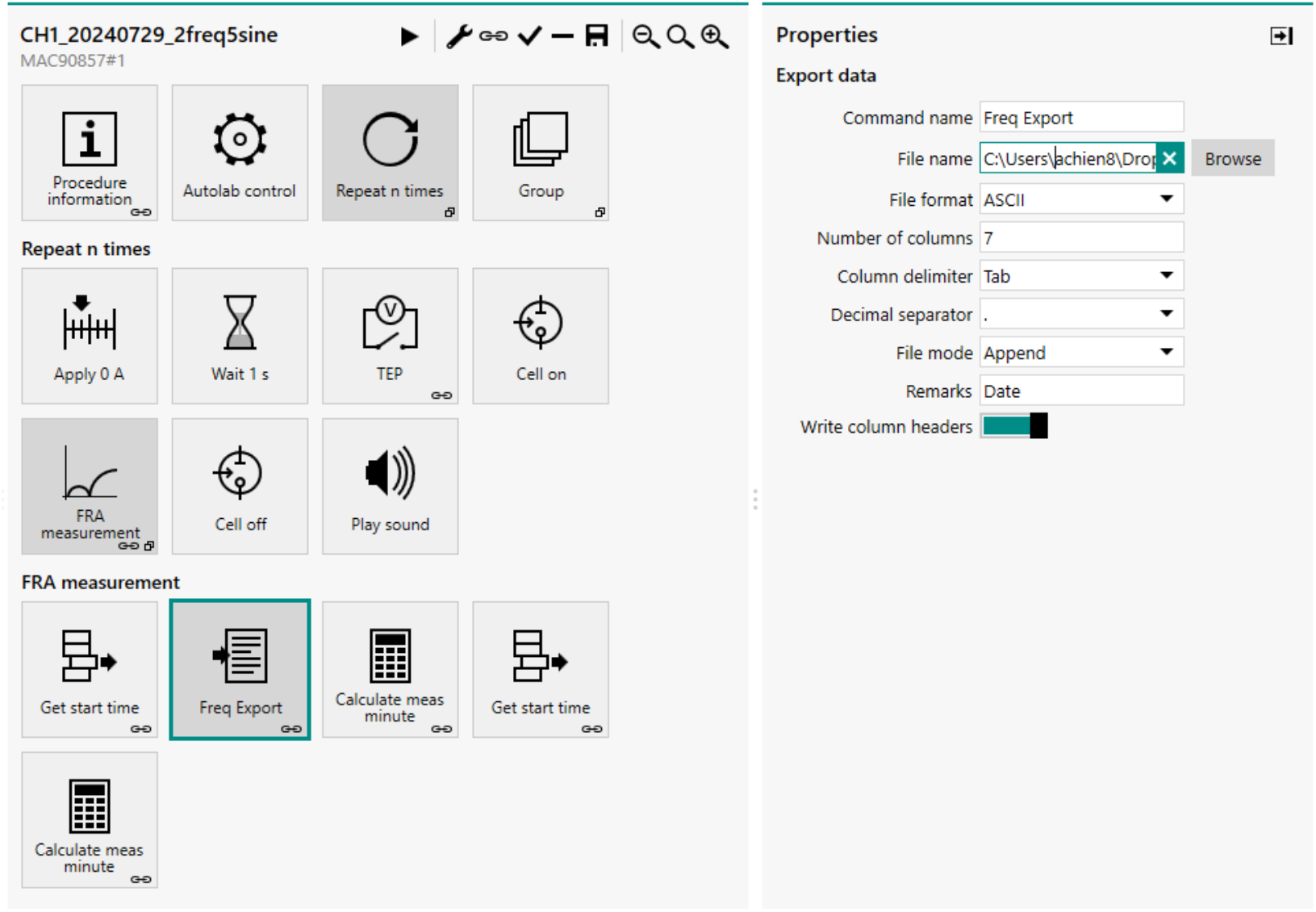
NOVA Protocol. Screenshot showing how to change filename on the right panel through the frequency export block.

**Figure 7.**
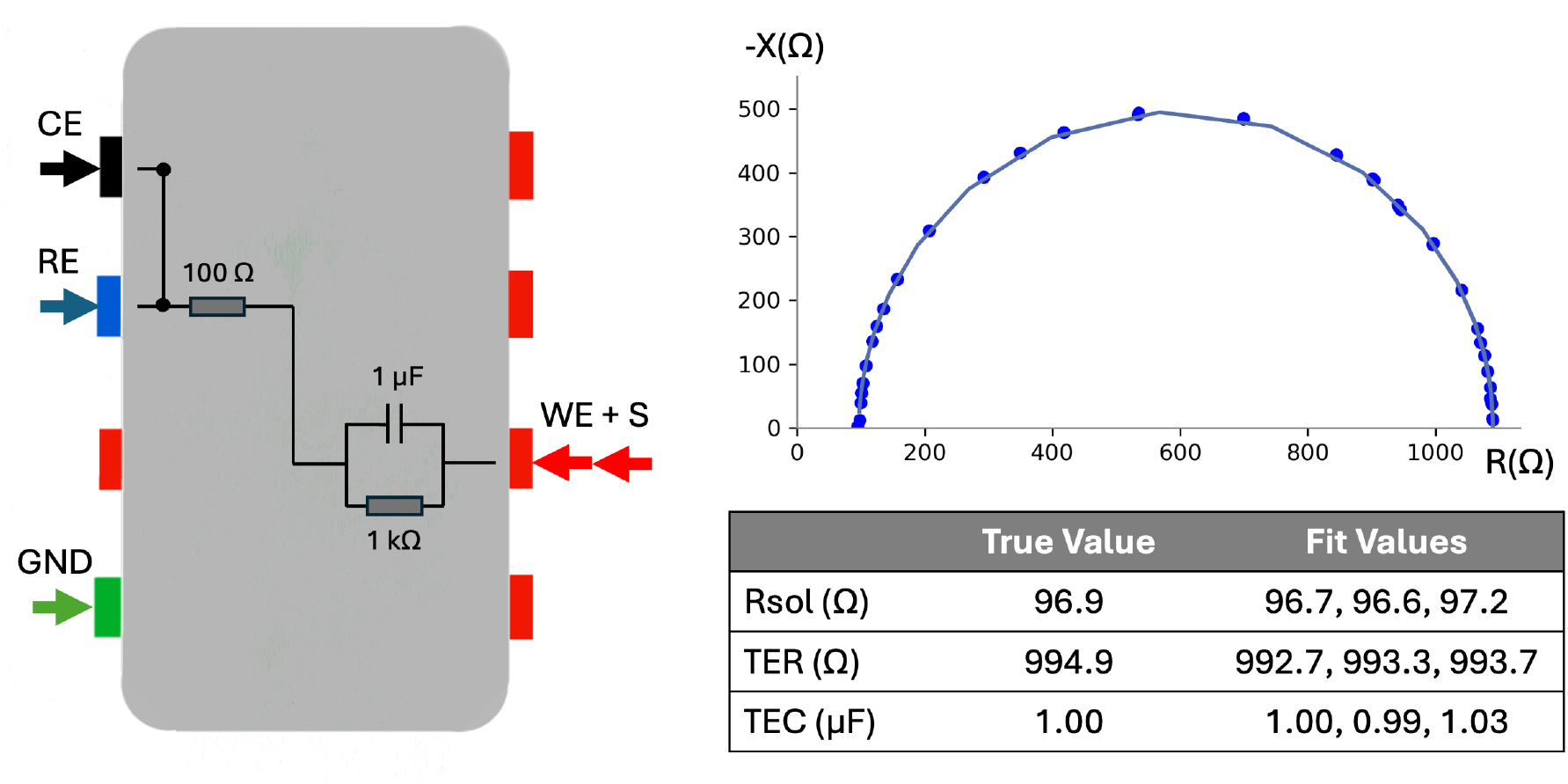
Model Cell Connections and Measurements. Three EIS measurements on test cell, 2 Hz - 50 kHz sweep, 2freq/dec, 5sines, connections to test cell shown on the left, Nyquist plot with raw data and fit lines shown on the right. Measurements and fit lines are indistinguishable between the three measurements, solution resistance and TER within 5 Ω, TEC within 0.03 *μ*F.

**Figure 8.**
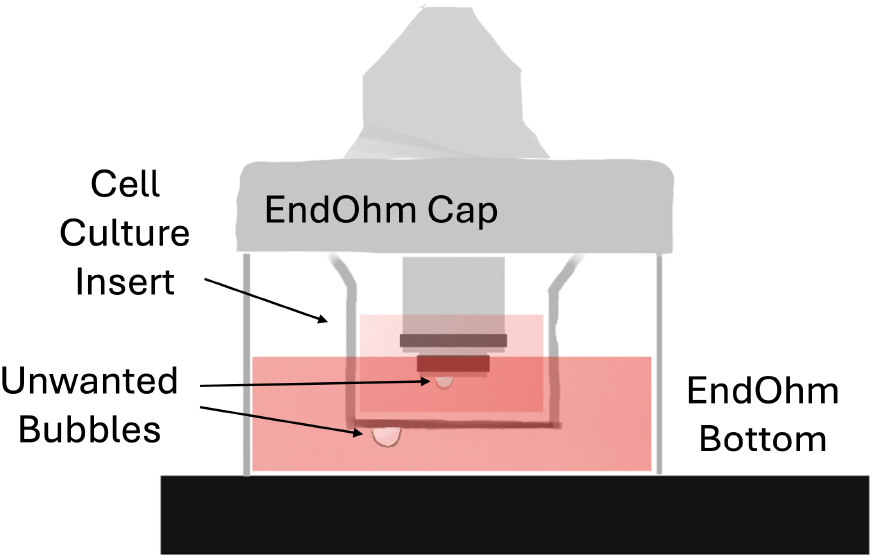
Bubbles in EndOhm Chamber. Bubbles easily form on the apical electrode surface and cell culture insert bottom surface upon submersion that contribute impedance and affect measurement accuracy. They should be prevented or removed as described in the procedure.

**Figure 9.**
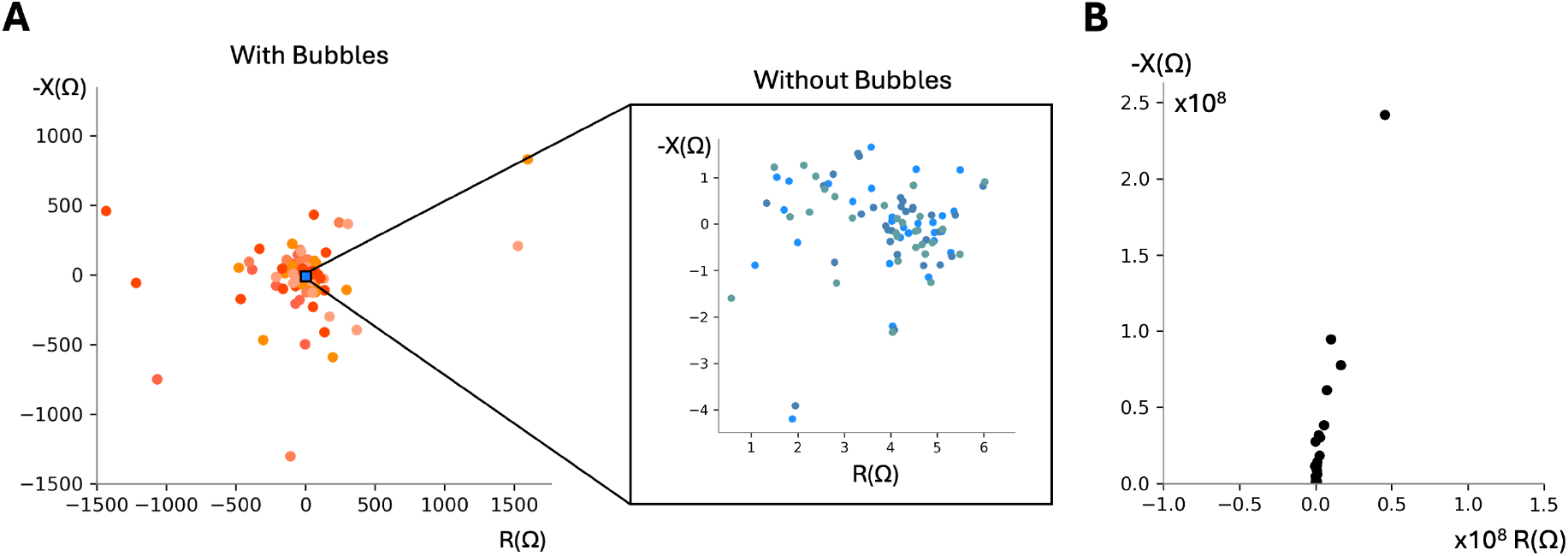
(A) Nyquist Plots of a Blank Cell Culture Insert With and Without Bubbles Present. Six representative noisy blank cell culture insert measurements due to bubbles each plotted in a different warm color, showing higher magnitude scatter. (Corning 3460 transwell in RPE media) (left). Three representative blank cell culture insert measurements without bubbles (right). All points remain within ±50Ω resistance and negative reactance for good measurements. Resistances can vary across inserts. We have observed resistance values clustered around a value from 0 − 50Ω resistance and 0Ω reactance. Note: We have set a threshold in the fitting code to switch to fitting only to a resistor (no capacitance) determined experimentally when the Nyquist plots look like this, determined experimentally when neither of the following statements are no longer true: the minimum reactance is less than three times the median reactance value measured for blank transwells and the sum of the *R*_1_ and *R*_2_ values calculated is greater than 9. (B) Nyquist Plot when apical electrodes are not submerged in apical solution gives resistance and reactance values on the order of 10^8^Ω and shape resembles slanted line. Frequency sweep 2 Hz - 50 kHz, 2freq/dec, 4 *μ*A current.

### 6.4 Epithelia Impedance Measurements

1. Begin epithelia impedance measurement either by continuing from the preceding steps 1-15 in order, or following the Experimental Setup section and then steps 1, 2, 4-5, and 9 in the Control Impedance Measurements section. The instrumentation should be connected as shown in Fig. 5.
2. Obtain confluent cells on cell culture inserts following protocols in [20] for bronchiolar or [21] for retinal epithelia.
3. Transfer cells from the incubator to the laminar flow hood and allow them to temperature stabilize for five minutes (within their existing well plate), cooling to approximately 26^°^C before beginning measurement.
4. While the cells are cooling, add 2.5 mL of the corresponding cell culture media to the EndOhm gently using the P1000 pipette and tips to avoid inducing bubbles.
5. Take out the sterile tweezers stored in reagent alcohol and shake lightly to dry off. Using the tweezers, place culture insert into the EndOhm bottom.
6. Place the EndOhm cap on top, lowering one edge in first to prevent bubbles. If bubbles appear, lift and reset the cap, or pipette media towards the bubbles to dislodge and pop them from the bottom of the EndOhm.
7. Change the file path and filename in the NOVA protocol to reference the cell culture insert as in Steps 6 and 7 of the Control Impedance Measurements Section (Fig. 6), and run measurement on the cell culture insert as in Step 8 of the Control Impedance Measurements Section. If the cells have grown confluent, the Nyquist plot should look like a semicircle or two adjacent semicircles intersecting. Representative examples of the Nyquist plot of bronchiolar epithelia is shown in Fig. 10A, showing the two semicircles intersecting as well as retinal pigment epithelia is shown in Fig. 10B.
8. Once the measurement is complete, take cell culture insert out of EndOhm, and replace with the next sample from the well plate. Basal media can remain in the EndOhm for all measurements on same cell replicates. All measurements from a single 12-well plate can be completed in 30 minutes.
9. When all measurements are complete, return cells to the incubator in their original well plate. Rinse EndOhm top and bottom with deionized water to remove proteins and charged particles from the surface of the electrodes, and let dry on the benchtop.

**Figure 10.**
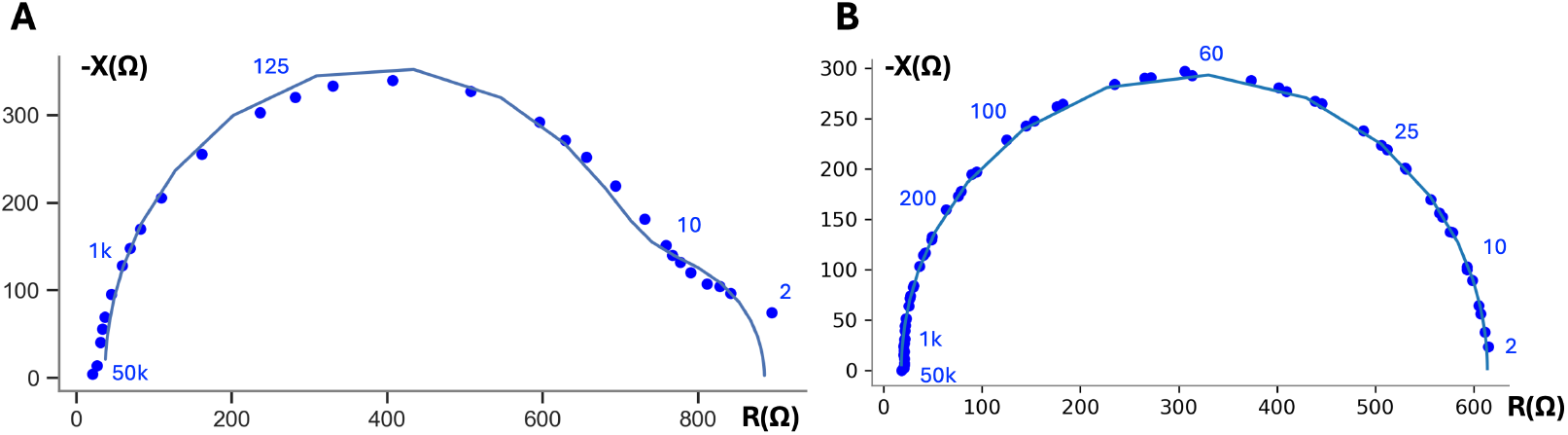
Representative Nyquist plot of 16HBE at day 7 (left) and mature RPE data (right) with frequency labels in blue, showing experimental data and fit line. Fit values normalized to 1.12 cm^2^ cross-sectional area: TER 950 Ω.cm^2^, TEC 1.06 *μ*F*/*cm^2^. Representative Nyquist plot of RPE (right), showing experimental data and fit line. Fit values normalized to 1.12 cm^2^ cross-sectional area: TER 667 Ω.cm^2^, TEC 4.10 *μ*F*/*cm^2^.

## 7. Data Analysis

In this section, we describe how the complex impedance data is analyzed to yield biological properties of interest: transepithelial resistance (TER), transepithelial capacitance (TEC), and membrane ratio. Specifically, we use a nonlinear least squares fitting algorithm, an established technique for fitting EIS data [23], to fit the impedance data collected in the preceding steps to the RCRC model in Fig. 1D.

A set of algorithms and steps are necessary to perform this analysis. For context, we first describe mathematically how this analysis is performed and then discuss how to quantify error.

### 7.1 Mathematical Model for Experimental Fit

Transepithelial impedance is complex and can be represented as the sum of two orthogonal values: resistance or real impedance, *R*(*ω*), and reactance or imaginary impedance, *X*(*ω*), where *ω* = 2*πf*, where *f* is the frequency in Hz, as shown in Fig. 9 and 10. This complex impedance, *Z*(*ω*), can be expressed as

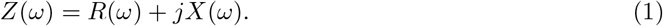

The measured complex impedance, *Z*, in Equation 1 can be mathematically modeled by combining the circuit elements in Fig. 1D such that

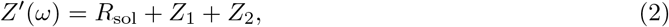

where Z_1_ is the impedance from the first parallel resistor-capacitor circuit and Z_2_ is the second as shown below.

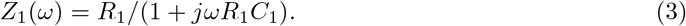

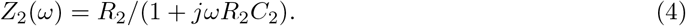

Note that *Z*^*′*^ is complex and can be separated into its real (resistive) and imaginary (reactive) components in a similar form to Equation 1:

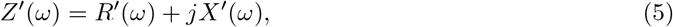

where

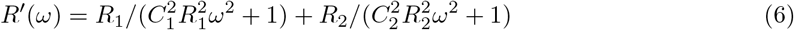

and

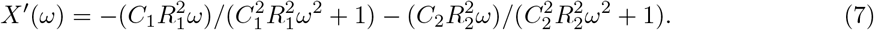

To relate *Z*^*′*^ (model) to *Z* (measurement), we use then use a nonlinear least squares solver to minimize an objective function, *J*, which is defined as the sum of the squared residuals, or error, between the measured and modeled data at all frequencies. J is calculated as

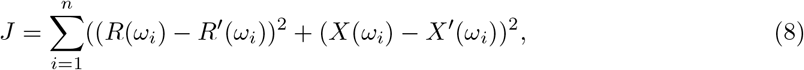

where *ω*_*i*_ represents each frequency. The values for *R*_sol_, *R*_1_, *R*_2_, *C*_1_, *C*_2_ that minimize the error are defined as the fit values.

In practice, we used Matlab’s nonlinear least squares solver to minimize *J* (Eqn 8). Specifically, the function ‘lsqcurvefit’ using code modified from Lewallen et al. 2023 [3] with the settings described in Table 1.

**Table 1:** Settings for nonlinear least squares fit. These non-default settings are configured in Matlab’s ‘optimoptions’ function and are applied to the ‘lsqcurvefit’ algorithm.

| Parameter | Value | Description |
| --- | --- | --- |
| MaxIterations | $3 \times 10^3$ | Maximum sets of parameter values ($R_s, R_1, R_2, C_1, C_2$) tried before reporting the best. |
| MaxFunctionEvaluations | $3 \times 10^3$ | Maximum number of function evaluations allowed. |
| FunctionTolerance | $3 \times 10^{-9}$ | Tolerance for convergence based on the function value. |
| OptimalityTolerance | $5 \times 10^{-7}$ | Tolerance for converging based on the gradient of the error function. |
| StepTolerance | $1 \times 10^{-9}$ | Tolerance for converging based on the step size of the error function. |

To mitigate the risk of converging to local minima or boundary values, each fitting procedure was initialized with a grid of 3 initial guesses per parameter. Resistance and capacitance values are limited to spanning the ranges of previously reported membrane properties in epithelial tissues [3, 10, 14, 19, 24–27] and adjusted based on observation:

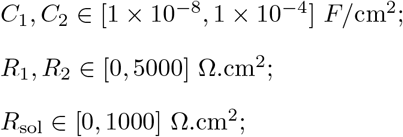

Following the minimization of the objective function *J*, the TER can be computed. TER is measured as the series combination of *R*_1_ and *R*_2_, then normalized by the cross-sectional area.

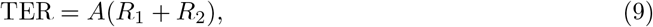

where *A* is the cross-sectional area of the tissue in cm^2^ and TEC is computed as the series combination of *C*_1_ and C_2_, also normalized to area:

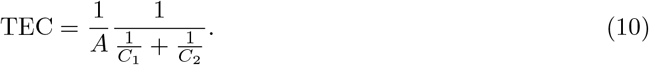

We also report the membrane ratio, *α*, as previously defined in Lewallen et al. [3]. For this circuit model, we define it as the ratio of the time constants of the two resistor-capacitor circuits as

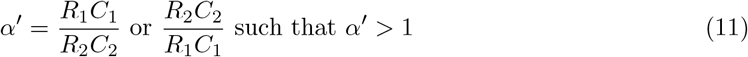

### 7.2 Data Normalization

Prior to fitting, we logarithmically scale and normalize the circuit element values, due to their significant difference in magnitude. This operation is reversed after fitting and is done to ensure the fitting algorithm assigns similar weight to the relative error of each parameter.

Specifically, two different normalizations are performed during each fit: (1) the magnitude of each circuit element, *R*_sol_, *R*_1_, *R*_2_, *C*_1_, *C*_2_, is converted to a log (base 10) domain, and (2) the real impedance and imaginary impedance are normalized to their respective maximum values. Impedance values were normalized by dividing each resistance value by its maximum resistance magnitude for that sweep, and dividing each reactance value by its maximum reactance magnitude for that sweep, such that the real and imaginary impedance all spanned the range of [-1, 1].

### 7.3 Error Quantification

Aside from the application cell electrophysiology, electrochemical impedance spectroscopy (EIS) is an established technique for studying electrode degradation and measuring battery properties. However, there are not well established metrics for quantifying error in fitting EIS data, and no studies on quantifying error for biological EIS data.

We have used the L_2_ norm of the residual (resnorm) in the past [3], as a reasonable error metric reported from Matlab’s nonlinear least squared fitting algorithm. Here, we further quantify how resnorm and other error metrics capture error differences in our biological data. We did a comparative study to determine which error metrics capture errors seen in biological samples best. We limited the study to modeling two different types of errors observed in our biological data: (1) random noise, modeled by Gaussian error, and (2) error at a single frequency. We began with six common error metrics used to quantify nonlinear fitting: residual norm, *χ*^2^, normalized *χ*^2^, mean absolute error (MAE), mean absolute percent error (MAPE), and root mean squared error (RMSE). Table 2 shows the equations used to calculate each of these metrics and the units in which they are reported.

**Table 2:**
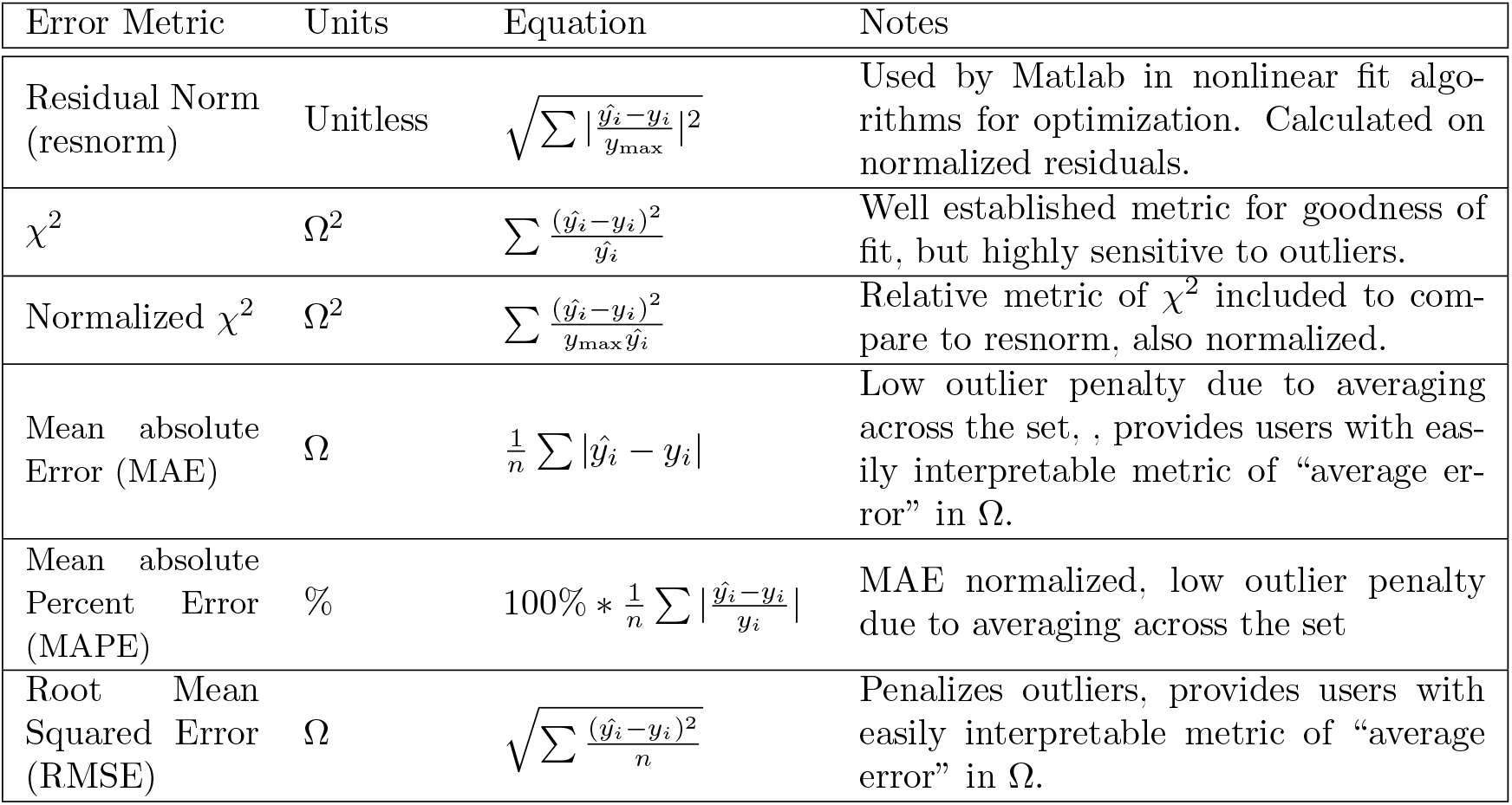
Error Metrics for nonlinear least squares analyses. Error metrics, equations, and descriptions. Each of these error metrics are calculated for the resistance and reactance, and then summed to output the single summary statistic for each sweep.

For modeling the first type of error, we added Gaussian noise to the reactance and resistance of the representative RPE data, to determine whether the error metrics can resolve differences between the original data, and data with errors sampled from the normal distributions *N* (0, 5^2^)Ω and *N* (0, 10^2^)Ω (Distributions in Fig. 11A, labelled 0 Ω, 5 Ω and 10 Ω). Representative Nyquist plots with the added error are shown in Fig. 11B. The resulting distributions of error for each error metric are shown as violin plots in Fig. 11C.

**Figure 11.**
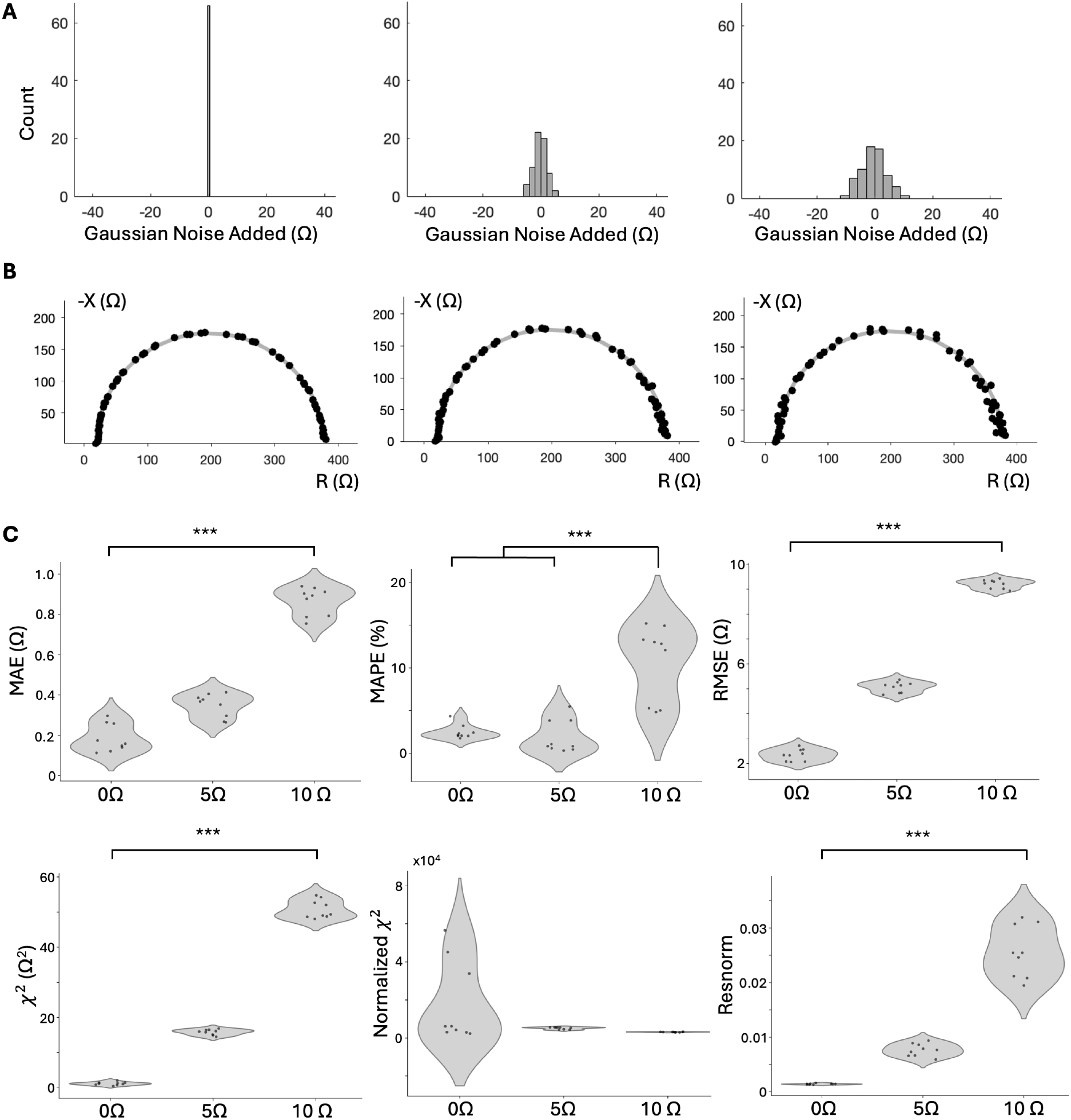
Error Comparison for Measurements with Gaussian Noise Added. (A) Error added to RPE data from Distributions *N* (0, 5^2^),*N* (0, 10^2^) (n=3 samples and technical replicates). (B) Representative Nyquist plots illustrating the varying noise added. (C) Violin plots showing mean absolute error (MAE), mean absolute percent error (MAPE), root mean squared error (RMSE), *χ*^2^, Normalized *χ*^2^, and resnorm metrics for original data, and data with 5 Ω and 10 Ω s.d. Gaussian noise added. ANOVA and Tukey HSD statistical tests were used to determine significance. (*** Tukey HSD p < 0.001, 1-way ANOVA *α <* 0.05, p < 0.01).

In good error metrics, we expect clear statistical differences across all the groups. Specifically, the error values from the 0 Ω group to be less than the 5 Ω group, which would be less than the 10 Ω group. We found MAE, RMSE, *χ*^2^ and resnorm show statistically significant differences and this increasing error trend between all three groups.

We also added single outliers to the data by adding 10 and 50 Ω noise to the resistance and reactance lowest frequency impedance point (2 Hz) and highest frequency impedance point (50 kHz) to assess which error metrics can capture the error due to single point outliers. The errors are shown in Fig. 12, where 0 Ω corresponds to the errors of the original data, LF stands for the low frequency value shifted (2 Hz), and HF stands for the high frequency value shifted (50 kHz), either 10 or 50 Ω, in both the positive resistance and negative reactance directions. Similar to adding Gaussian noise, we expect all of the error values from the original dataset to be shifted up from the original to the 10 Ω groups, and then again for the 50 Ω group. None of the metrics were sensitive enough to result in statistical significance between the 0 Ω and 10 Ω groups, but MAE, RMSE, *χ*^2^ and resnorm (similar to the Gaussian noise data) show statistical difference between the 10 Ω and 50 Ω groups. MAE and resnorm appear the best of these metrics because whether the low frequency or high frequency point is shifted, the violin plot shows the same distribution of error. As a result of these perturbations to the data, we concluded that MAE and resnorm both were able to capture both Gaussian and spurious noise as we defined it. Between MAE and resnorm, MAE is reported in Ω and is therefore more intuitive to understand. We concluded MAE should be used to assess the quality of the model fit, and report all fits (for RPE and 16HBE cells without noise added) have less than 5 Ω MAE. However resnorm is also a useful imbedded error metric reported in the summary table after fitting, while MAE requires running the data through an additional program.

**Figure 12.**
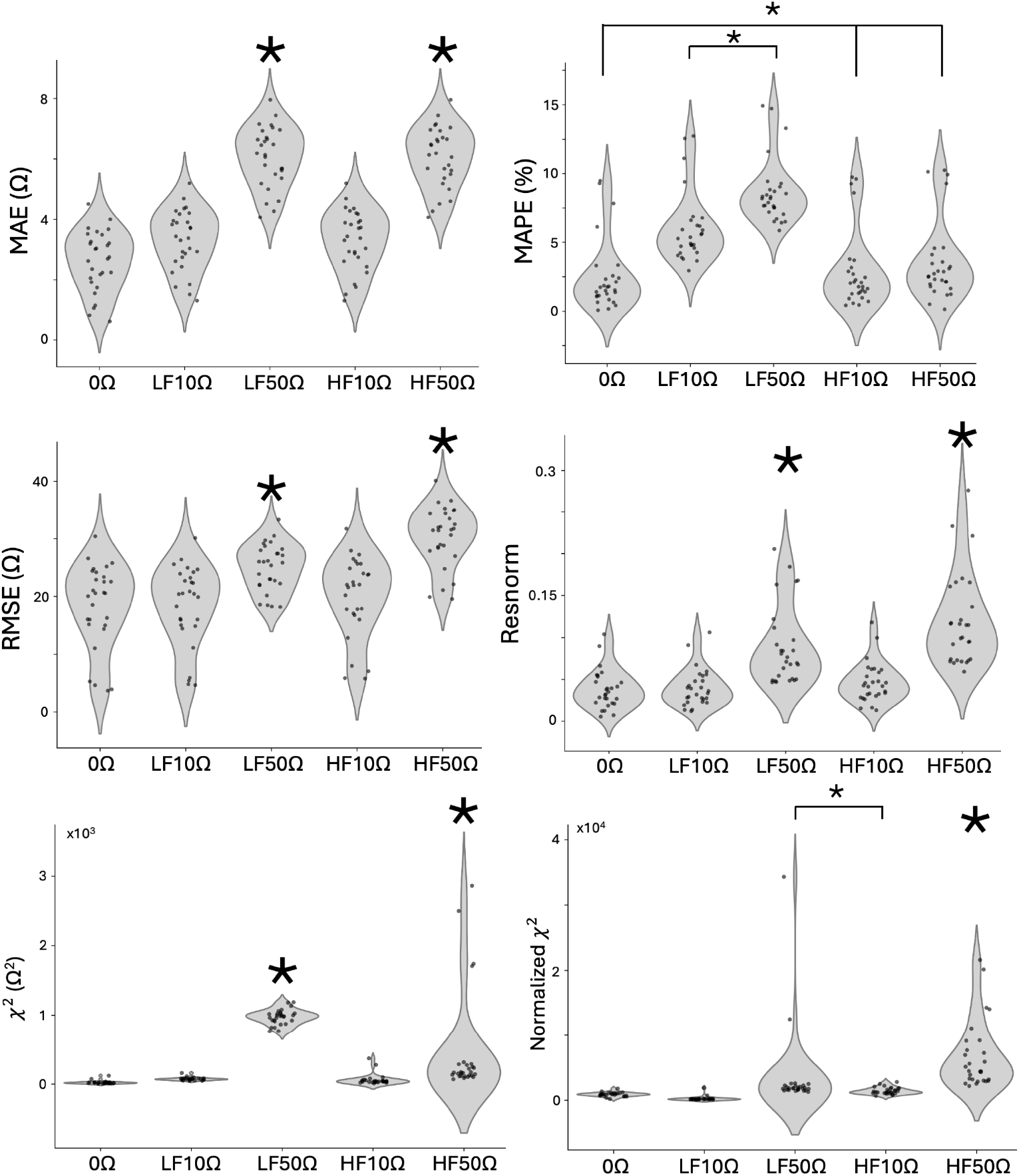
Error Comparison for Measurements with Single Frequency Error Added on data from four 16HBE samples day 3-9. 10 Ω and 50 Ω error added to resistance and reactance of 2 Hz (Low Frequency, LF) and 50 kHz (High Frequency, HF) 16HBE datapoints (n=28). Violin plots showing mean absolute error (MAE), mean absolute percent error (MAPE), root mean squared error (RMSE), resnorm, *χ*^2^, and Normalized *χ*^2^ metrics for original data, and data with 5 Ω and 10 Ω s.d. Gaussian noise added. 1-way ANOVA and Tukey HSD statistical tests were used to determine significance. Tukey HSD p < 0.001, ANOVA *α <* 0.05, p < 0.0001). (Groups with large * are statistically different from groups without, groups with small * mark statistical difference between two groups connected. e.g. MAPE * indicating LF10Ω and LF50Ω are statistically significant from the remaining groups.

### 7.4 Protocol for Data Analysis

To run the nonlinear least squares fit on the raw impedance data to calculate the TER, TEC, and membrane ratio (*α*), the impedance data files need to be copied into a specific folder, and a lookup table must be made with the cross-sectional area of each sample. Once run, the fitting code writes a summary table that outputs the TER, TEC, *α* and circuit fit values. Using the summary table and raw data, a Matlab script is then used to calculate the mean absolute error for assessing the accuracy of the fit values.

1. Open the the downloaded “extracellularEIS-main” folder from Github. Copy and paste the template lookup table in the “lookup table” folder titled “Template Lookup Table.xlsx” and rename the copy with your experiment name (e.g. “20240706Exp1 Lookup Table.xlsx”).
2. In the renamed lookup table, copy the name(s) of the raw data text file(s) (e.g. data1 freq.txt) you would like to fit into Column B, under “plateID.”
3. Type the cross-sectional area of the sample for each raw file in Column K, under “measArea” (1.12 cm^2^ for Corning 3460 cell culture inserts).
4. Number the files in column A with integer values starting at 1.
5. Save and close the lookup table.
6. Copy all the raw impedance data files output by the NOVA software into the “raw data” folder in the “extracellularEIS-main” folder.
7. Open the “NOVA batch 20240125.m” file in Matlab and change the file name and path in line 51 to point to the desired lookup table (e.g. “20240706Exp1 Lookup Table.xlsx”).
8. In the Matlab software, click the “Editor” menu in the top bar, and then the “Run” button to begin fitting. The software runs multiple fits in parallel, and should show “parallel processing” in the lower left corner. A figure will pop up and update with the raw data Nyquist plots and the overlaid fit line as the fit code is running through each sweep.
9. When completed, a file with the format “lookupTableName summary table.csv” will be generated in the “FIT” folder. The parameters output into the csv are listed in Table 3.
10. The mean absolute error (MAE) can then be calculated using the “calcMAE.m” program from the “otherCode” folder in the downloaded Github folder. Open the program in Matlab and change line 2 to reference the folder the raw data is located in. (e.g. dataLocation = ‘/Users/myuser/rawData/’;)
11. Change line 5 to reference the location of the summary table with the fit values. (e.g. SummaryTable = readtable(‘/Users/myuser/eisExtracellular/fit/20240706exp1 summary table.csv’);)
12. Click the “run” button in the top toolbar. The code writes a file titled “errors.txt” with the filenames in the first column, MAE in the second, and resnorm in the third column, written to same folder as the program.

**Table 3:**
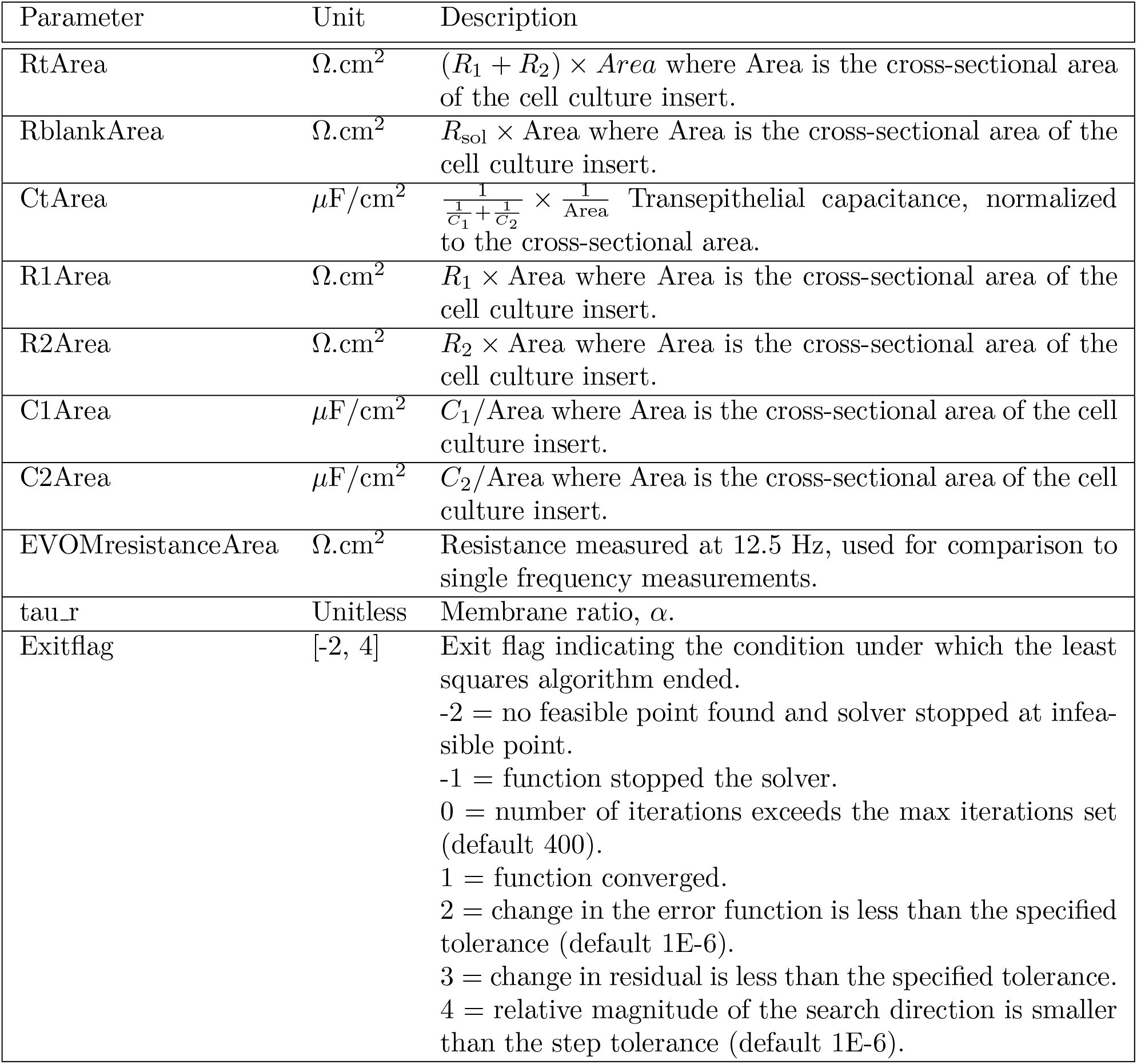
Fit Values Calculated by Nonlinear Least Squares Fitting, Output in Summary Table CSV File.

| Parameter | Unit | Description |
| --- | --- | --- |
| RtArea | $\Omega \cdot \text{cm}^2$ | $(R_1 + R_2) \times \text{Area}$ where Area is the cross-sectional area of the cell culture insert. |
| RblankArea | $\Omega \cdot \text{cm}^2$ | $R_{\text{sol}} \times \text{Area}$ where Area is the cross-sectional area of the cell culture insert. |
| CtArea | $\mu\text{F}/\text{cm}^2$ | $\frac{1}{\frac{1}{C_1} + \frac{1}{C_2}} \times \frac{1}{\text{Area}}$ Transepithelial capacitance, normalized to the cross-sectional area. |
| R1Area | $\Omega \cdot \text{cm}^2$ | $R_1 \times \text{Area}$ where Area is the cross-sectional area of the cell culture insert. |
| R2Area | $\Omega \cdot \text{cm}^2$ | $R_2 \times \text{Area}$ where Area is the cross-sectional area of the cell culture insert. |
| C1Area | $\mu\text{F}/\text{cm}^2$ | $C_1/\text{Area}$ where Area is the cross-sectional area of the cell culture insert. |
| C2Area | $\mu\text{F}/\text{cm}^2$ | $C_2/\text{Area}$ where Area is the cross-sectional area of the cell culture insert. |
| EVOMresistanceArea | $\Omega \cdot \text{cm}^2$ | Resistance measured at 12.5 Hz, used for comparison to single frequency measurements. |
| tau_r | Unitless | Membrane ratio, $\alpha$. |
| Exitflag | [-2, 4] | Exit flag indicating the condition under which the least squares algorithm ended.<br>-2 = no feasible point found and solver stopped at infeasible point.<br>-1 = function stopped the solver.<br>0 = number of iterations exceeds the max iterations set (default 400).<br>1 = function converged.<br>2 = change in the error function is less than the specified tolerance (default 1E-6).<br>3 = change in residual is less than the specified tolerance.<br>4 = relative magnitude of the search direction is smaller than the step tolerance (default 1E-6). |

## 8 Representative Epithelia Measurements

We applied the method as described to both RPE and 16HBE in order to provide examples of representative data from healthy cells. Fig. 13 shows the resulting TER, TEC, and membrane ratio, *α* on 3 samples of mature iPSC-RPE [1, 21], and 4 samples of 16HBE (cultured to maturity, measured on day 7 [28]). The RPE cells have an average TER of 587 Ω.cm^2^, an average TEC of 3.8 *μ*F*/*cm^2^, and average *α* of 2.0. The measurements a repeatable, with TER 500-667 Ω.cm^2^, and TEC 3.65-4.10 *μ*F*/*cm^2^. The 16HBE cells have an average TER of 983 Ω.cm^2^, an average TEC of 1.1 *μ*F*/*cm^2^, and average *α* of 20.0. The 16HBE measurements are also repeatable, with a narrower ranges for TER 955-1034 Ω.cm^2^, and TEC 1.07-1.10 *μ*F*/*cm^2^. These TER values are within the accepted threshold in literature for RPE [29, 30] and 16HBE [28]; other metrics of TEC and *α* have not been established.

**Figure 13.**
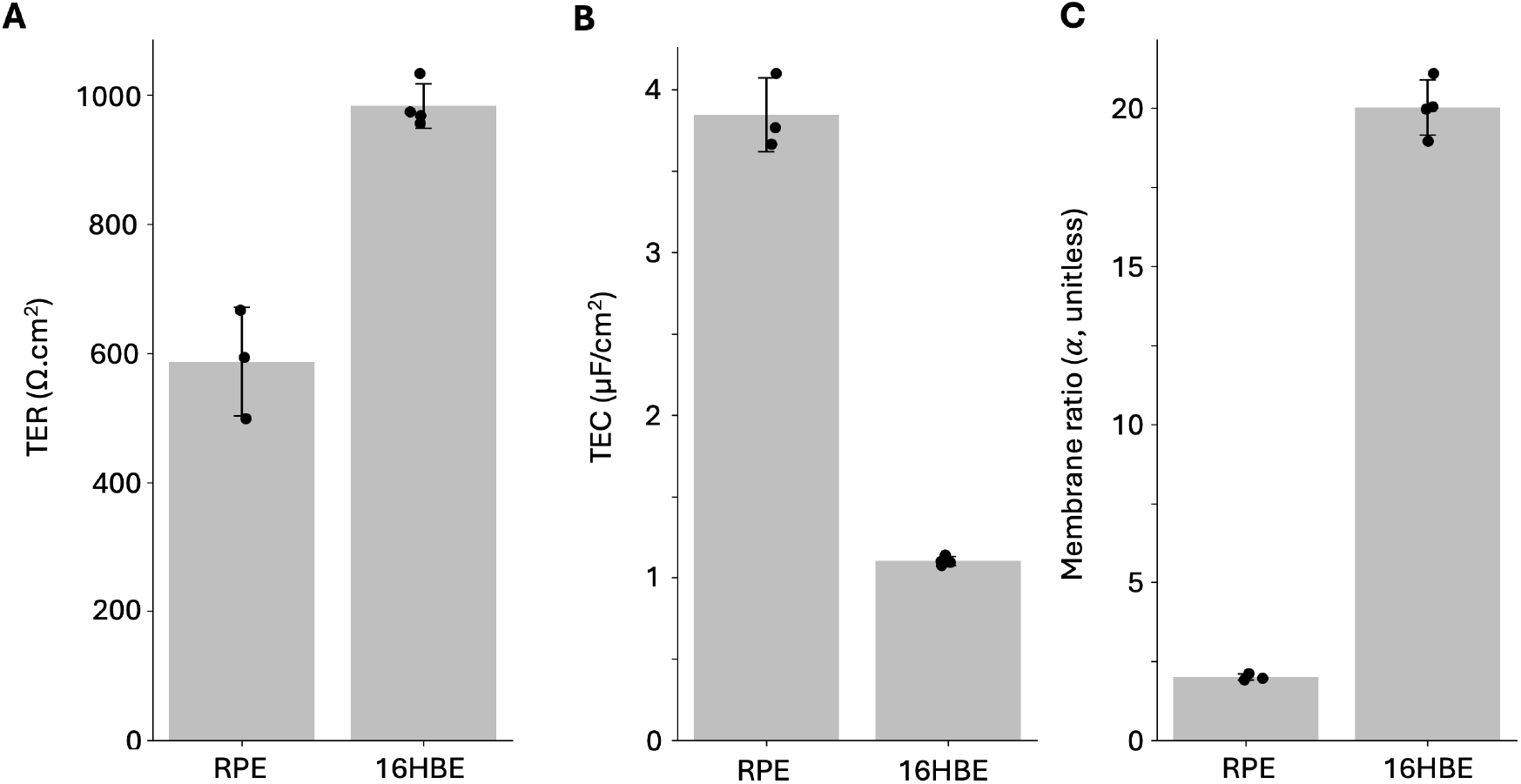
(A)TER, (B) TEC, and (C) membrane ratio measurements on mature RPE cells (n=3) and 16HBE cells (n=4, day 7) from frequency sweep 2 Hz - 50 kHz, 4 *μA* fit to RCRC model in Fig. 1D. Individual Nyquist plots are provided in the supplemental data, and fit values TER, TEC,*R*_sol_, *R*_1_, *R*_2_, *C*_1_, *C*_2_ are in the summary tables provided in the Github repository. Error values are in Fig. 11 and 12 (0 Ω groups). MAE was less than 6Ω for all 16HBE samples (n=4), and 1Ω for all RPE samples (n=3).

We have also used this technique to monitor EIS of bronchiolar epithelial cells, for example in a study of days 3-9 after seeding, reporting TER, TEC, and membrane ratios each day in Fig. 14. The TER trend shows the bronchiolar cells reach their highest TER by day 5 and maintain stable until day 8, while the TEC is stable around 1 *μ*F*/*cm^2^ days 5-9. The membrane ratio, similar to TEC, rises from day 3-5, before stabilizing around 20 through day 9.

**Figure 14.**
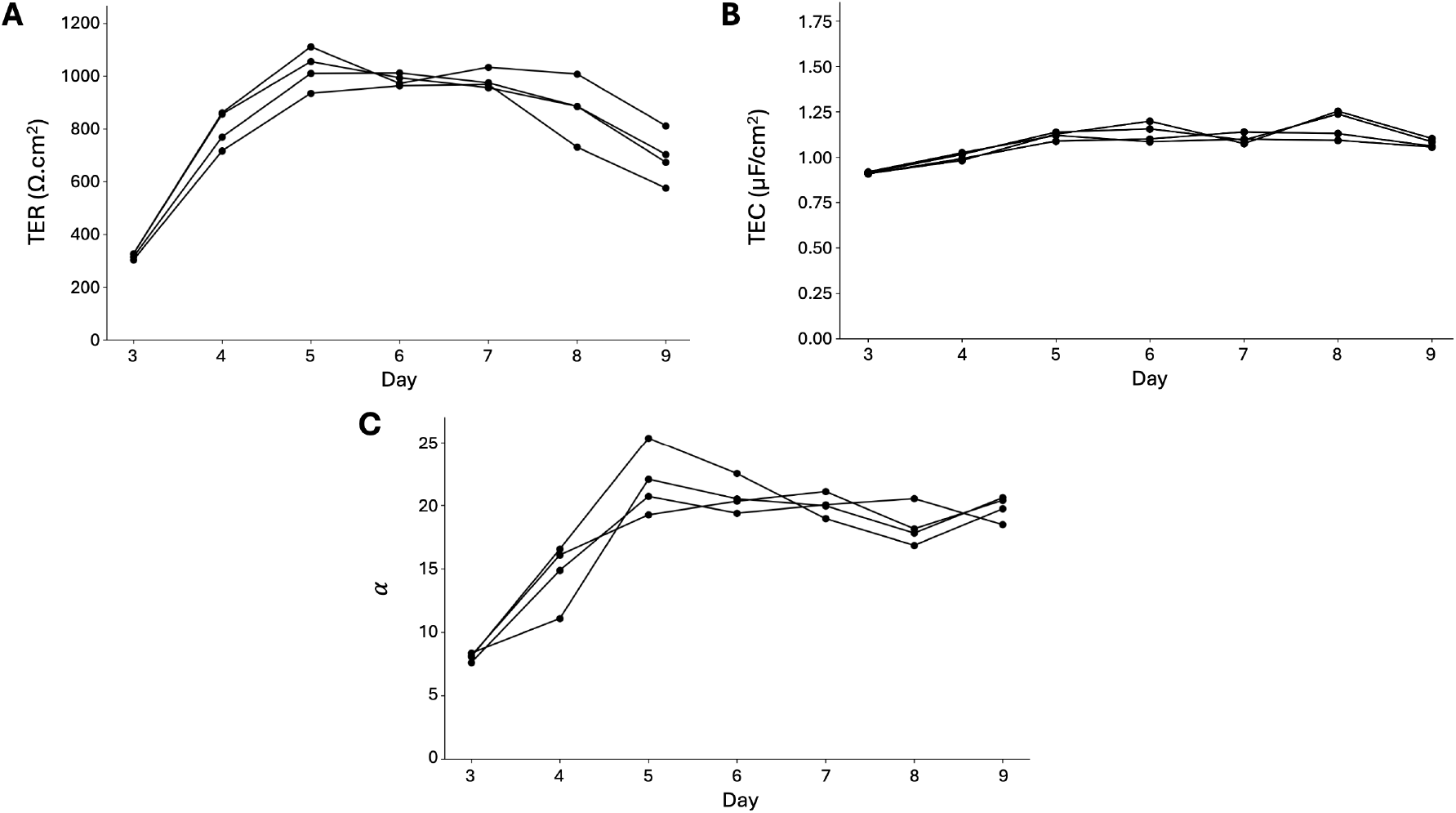
(A) TER, (B) TEC, and (C) Membrane ratio, *α* reported for 16HBE days 3-9 (n=4) from frequency sweep 2 Hz - 50 kHz, 5sines, 2 freq/dec., 4 *μA* fit to RCRC model in Fig. 1D. Individual Nyquist plots are provided in the supplemental data, and fit values TER, TEC,*R*_sol_, *R*_1_, *R*_2_, *C*_1_, *C*_2_ are in the summary tables provided in the Github repository. MAE was less than 10 for all samples.

## 9 Data Availability

All code and data associated with this protocol are openly available at https://github.com/chien0507/ extracellularEIS.git [DOI].

## 10 Notes

1. Is there a way to estimate TER and TEC directly from the Nyquist plot? Visually, the transepithelial resistance (TER) can be determined on the Nyquist plot as the difference between the two intersections with the real or resistance axis (848 − 37 = 811Ω normalized to the cross-sectional area 1.12 cm^2^ (908 Ω.cm^2^) as shown in Fig. 10A or 626 Ω.cm^2^ as shown in Fig. 10B). TEC for this RCRC model is more difficult to determine directly from the Nyquist plot, but each local maxima on the Nyquist plot can be used to determine the individual capacitances contributing if R_1_ and R_2_ are known, because *ωRC* = 1 at these points, as labelled in Fig. 2.
2. What is the expected exit flag for good fits? Typically the exitflag for good fits is 3, indicating the change in the residual is less than the specified tolerance, and the measurements of solution alone or a blank transwell are 1, indicating the function converged.
3. How does temperature affect TER? TER varies depending on the cell environment. Blume et al. discusses the variability in TER due to temperature [31]. Atmospheric changes due to changes in environmental conditions or solutions can also cause significant effects on the cells. The temperature dependence has only been reported on TER, but to try to measure as closely to physiologic conditions, in Ussing Chamber studies, cells are typically maintained at physiological temperatures. However one-time measurement of TER is commonly measured in the cell culture hood at room temperature because these conditions are easier to replicate, and relative changes in TER can still be reported and have biological significance.
4. How do media changes affect TER? We do not recommend changing media prior to measurement, as we have observed cells change impedance rapidly following a media change. Instead, change media after the measurement or several hours before the experiment to allow the cells to stabilize.
5. How frequently can you perform EIS in the EndOhm? Silver chloride electrodes like those used in the EndOhm are used for rapid signal transmission but are meant for short term measurements, as silver is a known toxin for cells. The chloride layer preventing such toxicity can easily be scraped off or nicked, having dose dependent toxic effects [32–34]. However exposure with smaller silver chloride electrodes has been used in alternative setups[35].
6. Are there known effects of these electrical measurement on cell behavior? Electrophysiology measurements for functional analyses can be non-invasive: the cells measured on whole cell culture inserts and only measured extracellularly (without penetrating the cells) can be returned to culture for later use and assessed multiple times. Generally, to minimize stimulation, magnitudes are kept below tens of microamps and several millivolts to prevent changes in activation of ion channels, cell shape, migration, and proliferation, although the exact values depend on the electrode type, culture media, and type of cells [34]. However, it is important to note that sending larger voltages or currents may influence the cell response, and are used to stimulate cells in conductive, inductive, or capacitive coupled systems [36]. For example, in bone marrow mesenchymal stromal cells, it has been shown that the membrane voltage can be manipulated to control proliferation using a DC signal of hundreds of millivolts [37].

## 11 Acknowledgements

Thanks to Analia Vazquez Cegla at Emory University in Prof. Nael McCarty’s laboratory for sharing cell culture protocols. Thanks to the research group of Dr. Kapil Bharti at the National Eye Institute within the National Institutes for Health for culturing and providing mature iPSC-RPE cells and media. OpenAI’s GPT-4 was used in the preparation of this document to generate initial drafts of the abstract and introduction sections that have since been further refined.

This material is based upon work supported by the National Science Foundation Graduate Research Fellowship under Grant No. (DGE-2039655). Any opinion, findings, and conclusions or recommendations expressed in this material are those of the authors(s) and do not necessarily reflect the views of the National Science Foundation. CRF acknowledges the NIH BRAIN Initiative Grant (NEI and NIMH 1-U01-MH106027-01), NIH R01NS102727, NIH Single Cell Grant 1 R01 EY023173, NIH R01DA029639 and NIH RF1AG079269, support from Georgia Tech through the Institute for Bioengineering and Biosciences, Invention Studio, and the George W. Woodruff School of Mechanical Engineering.

