## Supplemental Figure 15, Supplemental Figure 16, Supplemental Figure 17 for "Method for Extracellular Electrochemical Impedance Spectroscopy on Epithelia"

### 13 Supplemental Information

**Supplemental Figure 15: $R_{\text{sol}}, R_1, R_2, C_1, C_2$ values for electrical circuit control measurements on 4 test circuits modeling 16HBE and RPE observed impedances.**

| | Rsol | R1 | R2 | C1 | C2 |
|---|---|---|---|---|---|
| **16HBE1** | 27 Ω | 22 Ω | 266 Ω | 81.4 μF | 1 μF |
| **16HBE2** | 40 Ω | 300 Ω | 670 Ω | 49.0 μF | 1 μF |
| **RPE1** | 20 Ω | 20 Ω | 100 Ω | 83.7 μF | 1 μF |
| **RPE2** | 20 Ω | 83 Ω | 470 Ω | 82 μF | 5 μF |

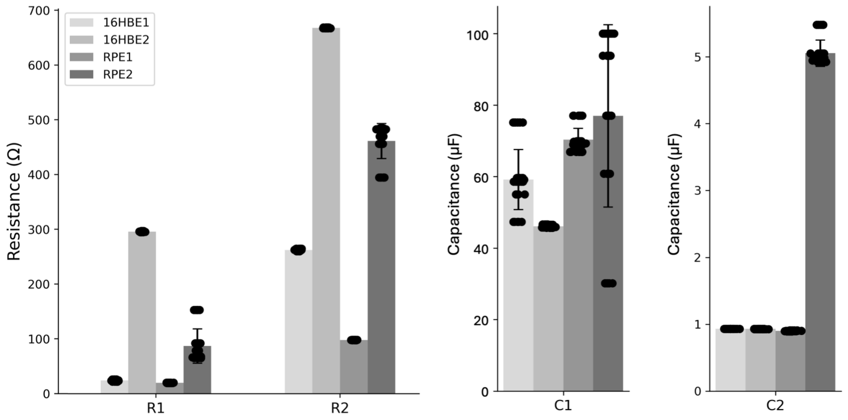


| | R1 error | R1 SD | R2 error | R2 SD | C1 error | C1 SD | C2 error | C2 SD |
|---|---|---|---|---|---|---|---|---|
| **16HBE1** | 2.24 | 1.32 | -3.61 | 1.49 | -22.3 | 8.40 | -0.07 | 0.00 |
| **16HBE2** | -4.07 | 0.64 | -2.07 | 0.71 | -2.9 | 0.38 | -0.07 | 0.00 |
| **RPE1** | -0.19 | 0.27 | -2.05 | 0.16 | -13.4 | 3.20 | -0.10 | 0.01 |
| **RPE2** | 4.16 | 31.28 | -8.54 | 31.91 | -5.0 | 25.52 | 0.05 | 0.20 |

Figure 15: $R_{\text{sol}}, R_1, R_2, C_1, C_2$ values for 4 test circuits, 6 technical replicates, 6 fits, showing the true values (top), bar chart of the raw values (middle), and error values (bottom). Error for RPE2 and 16HBE1 are worse, where membrane ratio is closer to 1, but still result in high precision and accuracy measurements of $\alpha$, TER, and TEC (Fig. 3). Resistor error and standard deviation in Ω, capacitance in $\mu$F.

**Supplemental Figure 16: Nyquist plots of 16HBE samples day 3-9.**

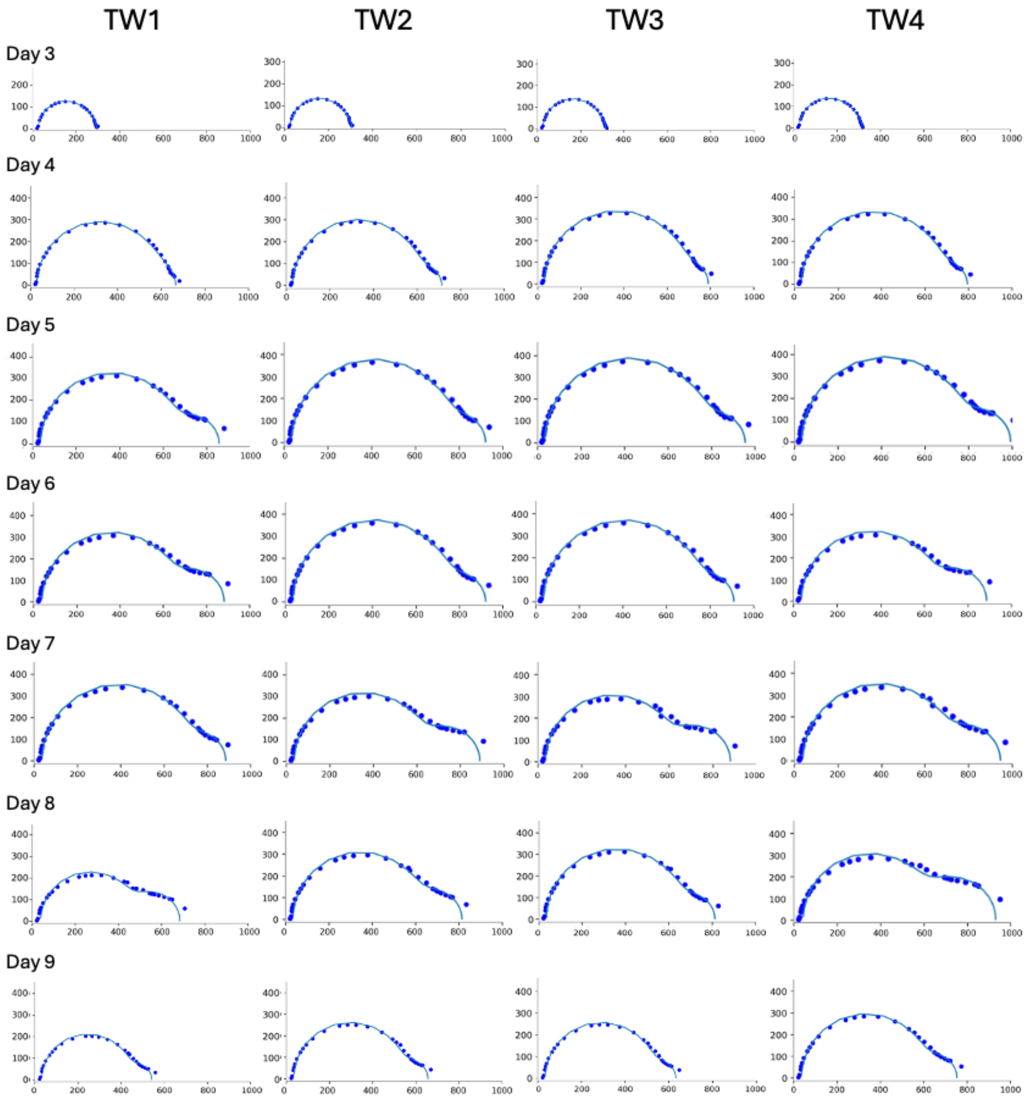


Figure 16: Nyquist plots (Y-axis negative reactance, x-axis resistance in Ω) for all four 16HBE samples from day three to day nine, with overlaid fit line. All four samples show largest growth of the Nyquist "bubble" occurs from day three to four, and significant shrinking by day nine.

**Supplemental Figure 17: Nyquist plots of three RPE samples with three technical replicates.**

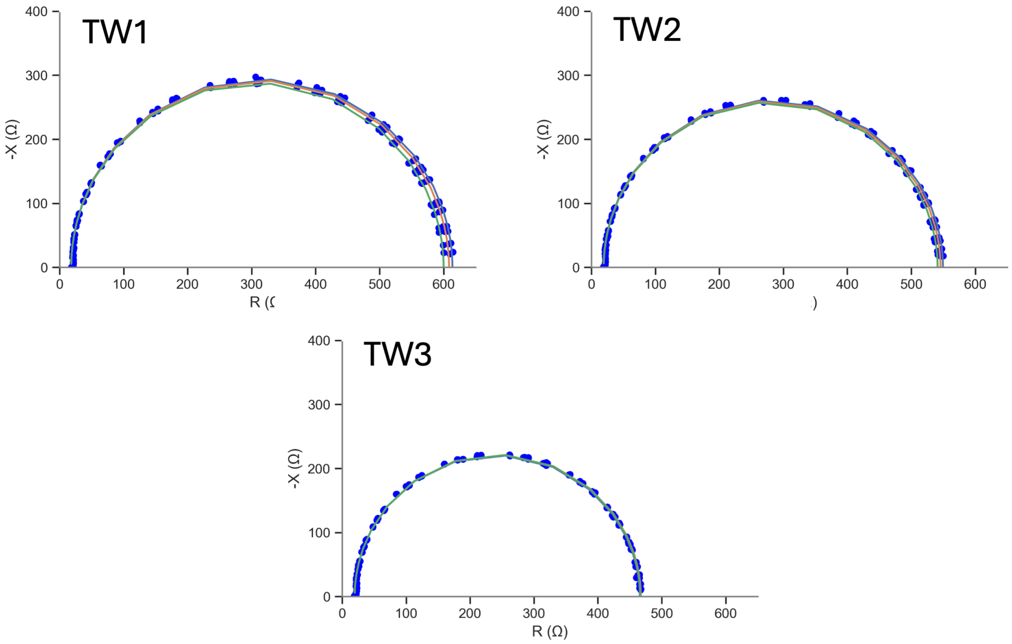


Figure 17: Nyquist plots (Y-axis negative reactance, x-axis resistance in Ω) for all six RPE samples, showing fit line and raw data. Little variability is observed across technical replicates, and fit values match Nyquist data well.
